# Novel exotic alleles of EARLY FLOWERING 3 determine plant development in barley

**DOI:** 10.1101/2022.07.15.500212

**Authors:** Tanja Zahn, Zihao Zhu, Niklas Ritoff, Jonathan Krapf, Astrid Junker, Thomas Altmann, Thomas Schmutzer, Christian Tüting, Panagiotis L. Kastritis, Steve Babben, Marcel Quint, Klaus Pillen, Andreas Maurer

## Abstract

*EARLY FLOWERING 3* (*ELF3*) is an important regulator of various physiological and developmental processes and hence may serve to improve plant adaptation which will be substantial for future plant breeding. To expand the limited knowledge on barley *ELF3* in determining agronomic traits, we conducted field studies with heterogeneous inbred families (HIFs) derived from selected lines of the wild barley nested association mapping population HEB-25. During two growing seasons, phenotypes of nearly isogenic HIF sister lines, segregating for exotic and cultivated alleles at the *ELF3* locus, were compared for ten developmental and yield-related traits. We determine novel exotic *ELF3* alleles and show that HIF lines, carrying the exotic *ELF3* allele, accelerated plant development compared to the cultivated *ELF3* allele, depending on the genetic background. Remarkably, the most extreme phenotypic effects could be attributed to one exotic *ELF3* allele, differing in only one SNP from the cultivated Barke *ELF3* allele. This SNP causes an amino acid change, which predictively has an impact on the protein structure of ELF3, thereby possibly affecting phase separation behaviour and nano-compartment formation of ELF3 and, potentially, also affecting its local cellular interactions causing significant trait differences between HIF sister lines.

## Introduction

Performance of crops like barley depends on their ability to adapt to different environments, which ultimately determines the yield potential. In context of an ever growing world population and climate change, maximizing crop yields for further food supply will be pivotal (FAOSTAT, 2009) and could be ensured, for example, by adaptation of crops to different environments (Challinor *et al*., 2014). More precisely, a meta-analysis of crop yield under climate change and adaptation based on 1,700 studies even predicted that cultivar adaptation would be the most promising way to increase yield under the predicted climate change (Challinor *et al*., 2014). Plant features important for plant adaptation are tolerance or resistance to abiotic and biotic stress factors like water and nutrient availability, extreme temperatures and soil salinity as well as pathogen infections. For a maximization of grain yield by adaptation, the exact timing of plant development and flowering time are particularly important (Alqudah and Schnurbusch, 2017; Cockram *et al*., 2007; Fernandez-Calleja *et al*., 2021; Francia *et al*., 2011; Wiegmann *et al*., 2019), for instance by adjusting the phenological stages to avoid periods of extreme stress (Cammarano *et al*., 2021; Kazan and Lyons, 2016; Zheng *et al*., 2013).

Flowering time is mainly controlled by day length and vernalisation (Andres and Coupland, 2012; Turner *et al*., 2005). To adjust flowering time, plants therefore need to be able to react to changes in photoperiod and temperature. For adaptation of barley cultivation to a wider range of environments, early flowering genotypes are necessary for short-growing seasons, while late flowering increases yield in temperate climates (Cockram *et al*., 2007; Fernandez-Calleja *et al*., 2021). The response to photoperiod under long day conditions in barley is mainly controlled by *PPD-H1*, a pseudo-response regulator, which promotes *VRN-H3*, a homologue of the *Arabidopsis thaliana* (Arabidopsis) *FLOWERING LOCUS T* (*FT*), through *CONSTANS* (*CO*), but also independently of *CO*, leading to the induction of flowering (Campoli *et al*., 2012a; Faure *et al*., 2012; Turner *et al*., 2005; Yan *et al*., 2006). Vernalisation requirement is mainly controlled by the interaction of *VRN-H1* (Yan *et al*., 2003), and *VRN-H2* (Yan *et al*., 2004), both affecting *VRN-H3*. While *VRN-H2* represses *VRN-H3*, *VRN-H1* is upregulated during vernalisation, leading to the activation of *VRN-H3* and repression of *VRN-H2* which in turn leads to the interruption of *VRN-H2* regulated *VRN-H3* repression, promoting the induction of flowering (Deng *et al*., 2015; Hemming *et al*., 2008; Yan *et al*., 2006). Due to a natural deletion of the entire *VRN-H2* gene, spring barley lacks the vernalisation requirement (Hemming *et al*., 2008; von Zitzewitz *et al*., 2005; Yan *et al*., 2004). Furthermore, there are genotypes that do not respond to photoperiod or vernalisation, making it possible to expand crop cultivation even further north. These genotypes have been characterized with *early maturity* (*eam*) or *earliness per se* (*eps*) loci (Faure *et al*., 2012). These loci may bring a new source of variation for the adaptation to different environments (Campoli and von Korff, 2014).

Flowering time is a complex trait which is controlled by a large regulatory network (Blümel *et al*., 2015). A central role in this network is taken by *EARLY FLOWERING 3* (*ELF3*), which is the focus of this study. ELF3 is an integral part of the circadian clock in both Arabidopsis and barley (Faure *et al*., 2012; Müller *et al*., 2020; Zagotta *et al*., 1996; Zakhrabekova *et al*., 2012). In general, the circadian clock is necessary to react and adapt to daily and seasonal environmental changes (Harmer, 2009; Wijnen and Young, 2006). It regulates a number of important genes that control plant growth processes and thereby contributes significantly to plant performance of important agronomic traits and adaptation to different environments (Bendix *et al*., 2015; Calixto *et al*., 2015; Nusinow *et al*., 2011).

The mechanistic understanding of the circadian clock is mainly based on studies in the model plant Arabidopsis, where *ELF3* functions as a core component of the clock (Nusinow *et al*., 2011; Thines and Harmon, 2010). Arabidopsis *ELF3* (*AtELF3*) is an oscillating gene with an expression peak in the early evening. *AtELF3* encodes a multifunctional protein that in turn regulates various physiological and developmental processes (Hicks *et al*., 2001; Nusinow *et al*., 2011), for example by repressing the activity of further core circadian clock genes (Dixon *et al*., 2011). Due to its diverse protein-protein interaction networking capabilities, AtELF3 presumably functions as a hub (Huang *et al*., 2016). Together with ELF4 and LUX ARRYTHMO (LUX), AtELF3 forms the evening complex (EC), a transcriptional regulator, which is an integral part of the circadian clock, repressing clock and growth-associated transcription factors (Huang and Nusinow, 2016; Nusinow *et al*., 2011). For loss-of-function *AtELF3* mutants an early flowering phenotype was shown (Zagotta *et al*., 1996) and in the context of this analysis it is important to note that AtELF3 controls photoperiod-responsive growth and flowering time, as well as temperature responsiveness of the circadian clock (Anwer *et al*., 2020; Jung *et al*., 2020; Zhu *et al*., 2022).

In barley (*Hordeum vulgare*), several clock orthologues from Arabidopsis have been identified with a high degree of conservation (Calixto *et al*., 2015; Campoli *et al*., 2012b; Müller *et al*., 2020). The gene *Praematurum-a* (*Mat-a*)/*EARLY MATURITY 8* (*EAM8*) was identified as a barley homologue of *AtELF3* (Faure *et al*., 2012; Zakhrabekova *et al*., 2012), from then on denoted as *HvELF3*. Its influence on flowering seems conserved since barley plants with a loss-of-function *elf3* also show early flowering phenotypes. Furthermore, those plants are insensitive to photoperiod and their circadian rhythm is disrupted (Boden *et al*., 2014; Faure *et al*., 2012; Zakhrabekova *et al*., 2012). Also, HvELF3 has recently been identified as a core component of the circadian oscillator since its absence leads to a non-rhythmic expression of other clock components (Müller *et al*., 2020), making it an essential regulator of the clock also in barley. Faure *et al*. (2012) have shown that *elf3* mutations lead to a higher expression of *PPD-H1*, particularly during the night, which subsequently induces *VRN-H3* and thereby earlier flowering. Also, under long day conditions, variation at *PPD-H1* was shown to influence flowering time of *elf3* mutants (Faure *et al*., 2012). Furthermore, in *elf3* mutants, altered expression of core clock and clock-output genes (*CO*, *VRN-H3, CIRCADIAN CLOCK ASSOCIATED1, GIGANTEA*, *TIMING OF CAB EXPRESSION1*) has been observed and increased expression of *HvFT1* (*VRN-H3*) was observed independently of *PPD-H1* (Boden *et al*., 2014; Faure *et al*., 2012). Ejaz and von Korff (2017) have shown later flowering under high ambient temperature for the cultivar Bowman, which harbours a functional *HvELF3* allele, whereas, for an introgression line with a non-functional *HvELF3* allele in a Bowman background, flowering time was accelerated. Furthermore, a larger reduction in floret and seed number has been observed for Bowman under high ambient temperature than for the introgression line. As such, *ELF3* (or natural variants/mutants thereof) contributed significantly to barley domestication and adaptation to higher latitudes by conferring a day-neutral flowering phenotype.

All barley research mentioned above is based on *elf3* loss-of-function mutants. We wanted to explore the role of natural barley *ELF3* variants, which is why we used the nested association mapping (NAM) population “Halle Exotic Barley” (HEB-25). The population originates from crosses of 25 highly divergent wild barley accessions (*Hordeum vulgare* ssp. *spontaneum* and *agriocrithon*, hereafter abbreviated as *Hsp*) with the elite cultivar Barke (*Hordeum vulgare* ssp. *vulgare*, hereafter abbreviated as *Hv*). The basis of our study were the results from a previous study on plant development traits in barley, where we identified a QTL region containing the *ELF3* locus (Maurer *et al*., 2016). This QTL significantly affected the traits shooting (SHO, Zadok’s stage Z31), shoot elongation phase (SEL, time from Z31 to Z49), heading (HEA, Z49), maturity (MAT, Z87) and plant height (HEI). Herzig *et al*. (2018) also describe the potential effects of *ELF3* on the traits SHO, HEA, MAT and HEI, confirming the results from the previous study in a different environmental context. In both studies the exotic *ELF3* alleles (*ELF3_Hsp_*) accelerated plant development and decreased plant height compared to the cultivated *ELF3* allele (*ELF3_Hv_*). Furthermore, *ELF3_Hsp_* effects for the mentioned traits varied for the 25 HEB families. Results for QTL1H10 (128-133.1 cM), the respective QTL for *ELF3*, were extracted from Herzig *et al*. (2018) and are shown in Fig. 1 for the trait HEA. Here, the exotic *ELF3_Hsp_* has varying effects among HEB families, but it is, in most cases, accelerating flowering (up to 2 days) compared to the cultivated barley *ELF3_Hv_* allele. Except for family 24, *ELF3_Hsp_* effects were stronger in Dundee (United Kingdom). Here, the maritime climate in 2014 and 2015 was characterized with colder summers, more and equally distributed rain and greater day lengths compared to Halle (Germany) with moderate-to-continental growing conditions (Herzig *et al*., 2018). The contrasting effects between the two field locations suggested that the *ELF3_Hsp_* effect on heading depends on environmental cues. Furthermore, Herzig *et al*. (2019) mentioned *ELF3_Hsp_* as potentially affecting grain nutrient content.

**Figure 1.**
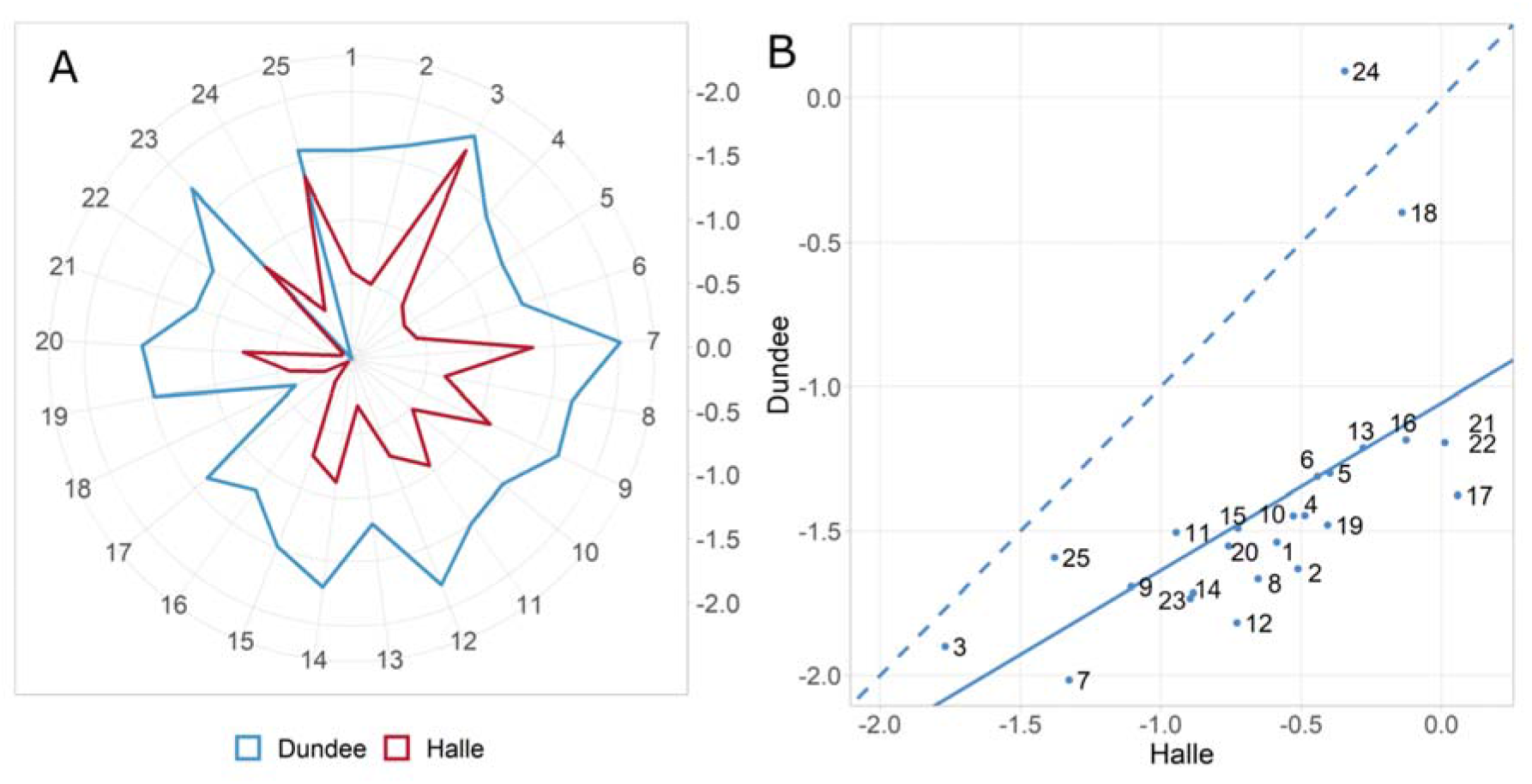
Family-specific effect diversity of exotic *ELF3* (*ELF3_Hsp_*) alleles compared to the cultivated *ELF3* (*ELF3_Hv_*) alleles for the trait heading in days. Comparison of all 25 families of the barley nested association mapping (NAM) population HEB-25 from Herzig *et al*. (2018). (A) Each line of the radar plot shows the respective HEB family with its average *ELF3_Hsp_* effect in days. (B) Scatterplot comparing *ELF3_Hsp_* flowering time effects in days between the two different locations Halle and Dundee. Regression line is shown as solid line and the dashed line is the diagonal separating effect strengths between the two locations. Pearson’s correlation coefficient between locations is 0.6.

However, despite these associations, there is little causal data about the effect of natural *ELF3* variants on barley flowering time regulation and crop performance. Xia *et al*. (2017) already described 51 *ELF3* haplotypes in 134 accessions (several cultivars, landraces and wild barleys, mainly native to the Qinghai-Tibet Plateau). Thereof, the novel *eam8.l* mutation was assumed to be responsible for an early flowering phenotype because of a mutation at position 3257 which probably contributed to intron retention and a truncated protein. However, other haplotypes also exhibited early flowering phenotypes, indicating the existence of other reasons for early flowering (Xia *et al*., 2017). As the selection of independent mutations at the *ELF3* locus might be a valuable tool to adapt barley cultivation to a wider range of environments (Faure *et al*., 2012), the aim of this study was to investigate the influence of further natural barley *ELF3* variants on several developmental and yield-related traits to subsequently identify *ELF3* alleles, which, in turn, may lead to an improvement of barley performance across environments. For this purpose, the barley NAM population HEB-25 was used as a basis for selection of Heterogeneous Inbred Families (HIFs). HIFs can be derived from advanced generations of lines with initial heterozygosity at a genomic region of interest. In this manner, allele effects can be estimated in a nearly isogenic background (Bergelson and Roux, 2010; Tuinstra *et al*., 1997). HEB-25 offers a diverse panel of wild barley alleles in a cultivated Barke background (Maurer *et al*., 2015). HIFs can be derived from those expected 6.25% of BC_1_S_3_ lines being heterozygous at *ELF3* to examine its association with a phenotype, enabling a direct comparison of allele effects on traits between two nearly-isogenic sister HIF lines segregating for the homozygous exotic and cultivated genotypes at *ELF3*.

Besides time to flowering (HEA), additional phenological traits were investigated such as time to shooting (SHO), duration of shoot elongation (SEL), duration of ripening phase (RIP, time from Z49 to Z87) and time to maturity (MAT). Furthermore, plant height (HEI), ears per square meter (EAR), grain number per ear (GNE), thousand grain weight (TGW) and grain yield (YLD) were investigated. Here, we describe significant effects of exotic *ELF3* variants on several agronomic performance traits, making *ELF3* an attractive target for future climate-resilient breeding approaches in barley. By investigating the *ELF3* coding sequences, we determine novel exotic *ELF3* alleles and show that an SNP in one exotic *ELF3* allele promotes expression of *VRN-H1*, potentially through a predicted altered protein structure of ELF3, which might reshape the phase separation behaviour and nano-compartment formation of ELF3.

## Materials and methods

### Plant material

For this study, HIFs were selected from the multiparental barley NAM population HEB-25 (Halle Exotic Barley; Maurer *et al*., 2015), which consists of 1,420 individual BC_1_S_3_ lines that were developed by an initial cross of the spring barley cultivar Barke (*Hordeum vulgare* ssp. *vulgare*) with 25 highly divergent wild barley accessions (*Hordeum vulgare* ssp. *spontaneum* and *agriocrithon*). For detailed information about the population design, see Maurer *et al*. (2015). In this study, HIF pairs were derived from HEB lines in generation BC_1_S_3:11_, which were heterozygous at *ELF3* in generation BC_1_S_3_. In addition, HIF pair 25_002_BC2 originates from a backcross of HEB line 25_002 (BC_1_S_3:7_), carrying the *ELF3_Hsp_* allele, with Barke. Here, the HIF pair was selected from the segregating progeny of the resulting BC_2_ plant. With the chosen plants (11 HIF pairs from 9 HEB families), two field trials were conducted in 2019 and 2020.

### Genotyping of HEB lines and HIF pairs

For a preselection of potential HIFs, existing Infinium iSelect 50k single nucleotide polymorphism (SNP) genotype data of HEB-25 was used (Bayer *et al*., 2017; Maurer and Pillen, 2019). Physical positions of SNPs were derived from the Morex reference sequence v2 (refseq2) (Monat *et al*., 2019). SNP data was first checked for quality, then an identity-by-state (IBS) matrix was created, coding homozygous Barke alleles as 0 and homozygous wild alleles as 2. Accordingly, heterozygous lines were coded as 1. Subsequently, the IBS matrix was converted to an identity-by-descent (IBD) matrix, as described in Maurer *et al*. (2017). This resulted in 32,995 SNP markers, which were used for preselection. Hereby, the first selection criterion was heterozygosity at the locus of interest, the *ELF3* gene (HORVU.MOREX.r2.1HG0078390.1). A gene specific marker (JHI_Hv50k_2016_57670), which is located inside *ELF3*, and flanking markers were used to determine whether heterozygosity was present at *ELF3* (Supplementary Table S1). Furthermore, lines showing heterozygosity at one of the other seven major flowering time loci in barley (Maurer *et al*., 2015) were discarded from the preselection, to ensure that no additional segregation in the background of *ELF3* would compromise the effect estimation of *ELF3* on traits, especially flowering time. Fifty plants of each BC_1_S_3:11_ line were grown and genotyped with kompetitive allele specific PCR (KASP) markers covering the *ELF3* region (Supplementary Table S2, Semagn *et al*., 2014) at TraitGenetics GmbH, Gatersleben, to select an *ELF3* HIF pair made of two nearly-isogenic lines segregating for *ELF3_Hv_* and *ELF3_Hsp_*. During the field trial 2019, the genotypes of selected HIF pairs were validated by TraitGenetics with the barley Infinium iSelect 50k chip (Bayer *et al*., 2017) and, subsequently, converted to an IBD matrix as described above (Supplementary Table S3).

### Field trials

In both years, 2019 and 2020, field trials were conducted at the ‘Kühnfeld Experimental Field Station’ of Martin Luther University Halle-Wittenberg (51°29′46.47″N; 11°59′41.81″E) to gather phenotypic data for 11 selected HIF pairs. Both field trials were sown in March (4^th^ of March 2019 and 19^th^ of March 2020), with fertilization and pest management carried out according to local practice. In 2019, the field trial was conducted in a randomized complete block design consisting of four blocks, each containing a randomized replication of the 11 selected HIF pairs. The plots consisted of 3 rows (50 seeds each) with a length of 1.5 m and a distance of 0.15 m between rows. Plots were evenly spaced by 0.3 m. The field trial in 2020 was conducted in six randomized blocks. The plots consisted of 8 rows with a length of 3.2 m, a distance of 0.15 m between rows, 0.3 m between plots and a seeding density of 300 seeds per m^2^. Both sister lines of a HIF pair were always sown next to each other for comparison and to minimize spatial effects. Additionally, elite donor Barke was placed as a control in 27 plots in 2019 and 11 plots in 2020.

### Environments

The growth period had the same length in both years, only in 2020, sowing was carried out two weeks later than in 2019 (4^th^ of March 2019 and 19^th^ of March 2020) as earlier sowing was not possible due to too much rain and wet soil in 2020. Therefore, maturity of the latest line was two weeks earlier in 2019. During the respective growth periods of the field trials, the mean temperature was 0.5 °C higher in 2020 (13.4 °C), especially in the third month of the vegetation period when heading started, temperature was on average 3 °C higher in 2020. However, during the last month of the respective growth period, temperature was almost 2 °C higher in 2019 (21.3 °C) with high daily average temperatures of up to 29 °C. In 2020, the highest daily average temperature was 23.8 °C (Supplementary Fig. S1, Supplementary Table S4). The sum of precipitation over the whole growth period was almost the same in both years (approx. 127 mm). While in 2019 rainfalls occurred during spring, directly after sowing and equally distributed over the summer, almost no rain occurred during the first third of the vegetation period and almost 50% of rain during the last month of the vegetation period in 2020 (Supplementary Fig. S1, Supplementary Table S4). Due to later sowing in 2020, the day was one hour longer in the beginning of the experiment in 2020 and the longest day (21^st^ of June) was later in the vegetation period in 2019 than in 2020, leading to a larger absolute amount of daylight in 2020 (cumulative day length (CDL): 1775:42 h in 2020 compared to 1715:42 h in 2019, Supplementary Table S4).

### Phenotypic data

Phenotypic data were recorded in both years for ten developmental and yield-related traits (Table 1). For the developmental traits SHO until MAT, growing degree days (GDD) were calculated following equation (1) of McMaster and Wilhelm (1997) with a base temperature of 0 °C. The decision, for which trait days or GDD was used, is based on estimated repeatabilities (Rep) and heritabilities (H^2^) (Supplementary Note S1, Supplementary Table S5).

**Table 1.**
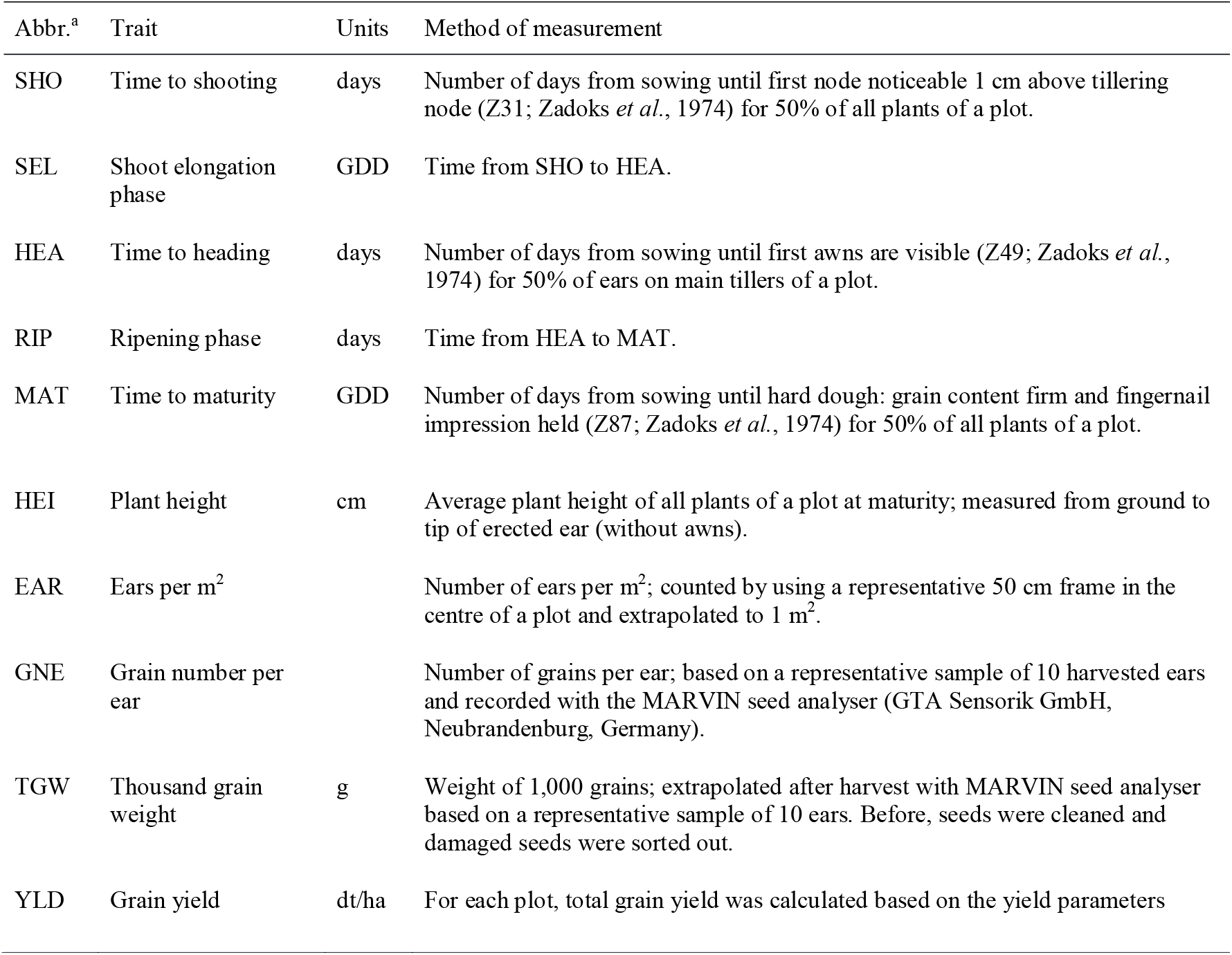
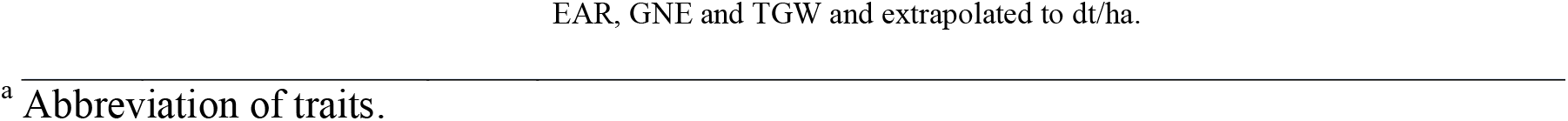
List of evaluated traits.

### Image-based phenotyping in controlled environments

To validate barley *ELF3* effects in a controlled environment, a phenotyping experiment was conducted using the LemnaTec system at the Leibniz Institute of Plant Genetics and Crop Plant Research (IPK) in Gatersleben. One HIF pair (10_190), the cultivar Bowman and two *elf3* mutants in a Bowman background (BW289 and BW290, carrying the *eam8.k* and *eam8.w* alleles, respectively (Faure *et al*., 2012; Zakhrabekova *et al*., 2012)), were grown in 13-15 replications per genotype for 64 days (sowing on November 18^th^ 2019) under standard conditions with day/night temperatures of 20 °C/18 °C and long days (LD) with 16 h light and 8 h darkness, respectively. Top- and side-view RGB images (LemnaTec automatic phenotyping system at IPK Gatersleben) from each plant were taken every day after day 8 and every two to four days after day 33. All analysed growth and developmental parameters were scored from these images. To obtain data for plant height, area and volume, the Integrated Analysis Platform (IAP) pipeline was used (Klukas *et al*., 2014). Number of tillers was counted manually on day 64 (all tillers were included). On day 64, the aerial parts of the plants were harvested, and fresh weight was measured using an electronic scale. Dry weight was measured after placing plant material into a drying oven for 3 days at 80 °C.

### Statistical analyses

All statistical analyses were performed either with SAS 9.4 (SAS Institute Inc., Cary, NC, USA) or R 3.4.3 (R Development Core Team, Vienna, Austria). Basic descriptive statistics and comparative statistics between HIF sister lines were calculated in R using the compare_means method ANOVA. SAS PROC HPMIXED was used to estimate best linear unbiased estimators (BLUEs), assuming fixed genotype and block effects in a linear mixed model. Pearson’s correlation coefficients were calculated using the CORRGRAM package in R. BLUEs were used for calculation of correlation of traits across years and individual values were used for correlation of traits within a year.

Repeatabilities (Rep) for each year and broad-sense heritabilities (H^2^) were calculated as

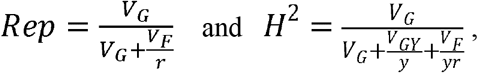

where V_G_, V_GY_ and V_F_ correspond to the genotype, genotype × year and error variance components, respectively. The terms y and r represent the number of years and replicates, respectively. For estimation of variance components with SAS procedure PROC VARCOMP, all effects were assumed to be random. Furthermore, an ANOVA was conducted for each trait to test for significant genotype and year effects as well as for significant genotype × year interactions. For the image-based trial, ANOVA was conducted pairwise with SAS to test for significant trait effects between Bowman and the two mutants as well as between the two HIF sister lines of 10_190 for plant height, area and volume. For the traits heading, number of tillers, fresh weight and dry weight, ANOVA was conducted to test for significant differences between all five genotypes (pairwise).

### ELF3 coding and protein sequence

The full-length genomic sequence of *ELF3*, from original wild barley donors, Barke, Bowman and BW290, was amplified using Ex Taq DNA Polymerase (Takara Bio, Kusatsu, Shiga, Japan). The purified amplicons were submitted to Eurofins Genomics (Ebersberg, Germany) for dideoxy sequencing. Five to six overlapping fragments were assembled, and the coding sequence was then obtained by alignment with the reported *ELF3* sequence from cultivar Igri (GeneBank accession number HQ850272; Faure *et al*., 2012). Subsequently, the protein sequences were obtained by using ExPASy translate tool (Gasteiger *et al*., 2003). In addition, the promotor region of *ELF3* in HIF sister lines of 10_190 was sequenced. Therefore, genomic DNA was amplified using ALLin™ RPH Polymerase from highQu (https://www.highqu.com/ALLin-RPH-Polymerase/HLE0101) and purified via Thermo Scientific™ GeneJET PCR Purification Kit (https://www.thermofisher.com/order/catalog/product/K0701) before it was Sanger sequenced by Eurofins Genomics (Ebersberg, Germany). Primers used for PCR and sequencing are given in Supplementary Table S6.

Structure of Barke *HvELF3* was visualized by using Exon-Intron Graphic Maker (Bhatla, 2012) available at http://wormweb.org/exonintron (accessed November 29, 2021). AtELF3 protein (Col-0) was obtained at https://www.arabidopsis.org/ (AT2G25930, The Arabidopsis Information Resource (TAIR), accessed November 29, 2021) and three AtELF3 protein domains were defined according to Nieto *et al*. (2015). For alignment with AtELF3, Barke ELF3 was obtained as described above and Morex ELF3 sequence from Morex reference sequence v2 (Monat *et al*., 2019). Alignment of AtELF3 and HvELF3 (of Barke/Morex) sequences was done using MAFFT version 7 (Katoh *et al*., 2019; Kuraku *et al*., 2013) available at https://mafft.cbrc.jp/alignment/server/ (accessed November 29, 2021) and, subsequently, the respective HvELF3 protein domains were retrieved.

Furthermore, for comparison purposes of the coding sequence variation in the HEB-25 wild donors with already described variation in the literature, *ELF3* coding sequences were extracted from Xia *et al*. (2017) as well as from NCBI (https://www.ncbi.nlm.nih.gov/, accessed May 4, 2022) and Jayakodi *et al*. (2020), using the IPK Galaxy Blast suite (https://galaxy-web.ipk-gatersleben.de/, accessed May 4, 2022). Determination of *ELF3* haplotypes based on exon sequences was done using the “haplotypes” package in R.

### Gene expression analysis

To test for differences in gene expression between HIF sister lines, HIF pairs 10_003 and 10_190 as well as plants of Bowman and BW290 were grown for 16 days under LD with 16 h light and 8 h darkness, day/night temperatures of 20 °C/18 °C, and light intensity of 300 μmol m^−2^s^−1^. On day 17, starting from the onset of light (ZT00), leaf samples were harvested every 4h, using three biological replicates. Total RNA was isolated using the NucleoSpin RNA Plant Kit (Macherey-Nagel), cDNA was synthesized using the PrimeScript RT Reagent Kit (Perfect Real Time, Takara Bio), and quantitative real-time-PCRs were performed on an AriaMx Real-Time PCR System (Agilent) using Absolute Blue Low Rox Mix (Thermo Fisher Scientific). The reference genes *HvGAPDH* and *PTF* were used for normalization. Primer sequences are described in Supplementary Table S6. Differences between Bowman and BW290, or between HIF sister lines at each time point were analyzed for significance by a two-sided Student’s *t*-test using GraphPad QuickCalcs (http://graphpad.com/quickcalcs/).

### ELF3 protein sequence structure analysis

For global structure prediction a local installation of Alphafold v2.1.0 (Jumper *et al*., 2021) with max_template_date=2021-10-12 and model_preset=monomer_ptm was used. Results were analysed by an in-house python script (available upon request).

To identify structural homologues, the BLASTp webserver with the database “Protein Data Bank proteins (pdb)” and the ELF3 sub-sequences “SSRGSELQWSSAASSPFDRQ” and “SSRGSELQGSSAASSPFDRQ” were used. The derived hits were analysed by an in-house PyMOL script (available upon request) regarding their structural completeness (min. 5 resolved residues in the pdb file) and their annotated secondary structure. The weblogo was generated using the Berkley weblogo webserver (Crooks *et al*., 2004).

For disorder analysis, the amino acid sequences of the barley homologues from the annotated Arabidopsis proteins were identified using BLASTp. A local installation of the MobiDB-lite suite (Necci *et al*., 2017) was used to predict the disorder content of the derived sequences.

## Results and discussion

### Phenotypic variation suggests year-by-year analysis

We observed broad variation for all traits, both between genotypes and years (Supplementary Fig. S2, Tables S7 and S8) with medium high coefficients of variation (CV) in both years. As expected, for the elite parent and control cultivar Barke, the CV was not as high as the CV across the studied HIF pairs. CVs for YLD were particularly high, which can be explained by the high variation of EAR. An ANOVA revealed significant (p < 0.001) effects for genotype and year as well as for genotype × year interaction except for SEL, MAT and TGW between the two years and RIP, EAR, TGW and YLD for genotype × year interaction (Supplementary Table S9). In 2020, for all developmental traits, except RIP and SEL [GDD], plants showed a faster development than in 2019 (Supplementary Table S8). For the trait SEL, plants showed a faster development in 2020 when comparing this growth phase in days, while GDD values were lower in 2019, showing that the average temperature in 2020 was higher during this growth period than in 2019. Furthermore, plants were smaller in 2020 and all yield components had lower values. Especially, yield was unexpectedly low in 2020. Presumably, the generally faster phenological development in 2020 led to a shorter growth period (e.g. due to different weather conditions, Supplementary Fig. S1, Table S4) and left the plants less time for assimilation, grain filling and biomass production, resulting in smaller plants, lower yield components and, consequently, lower grain yield. The average grain yield for spring barley in Germany in 2020 was 55.6 dt/ha (Federal Ministry of Food and Agriculture, 2020). In this study, YLD was very high in 2019 (97.6 dt/ha) whereas in 2020 it was far below (38.8 dt/ha). Due to the small plot size in 2019, yield was probably overestimated. Repeatabilities (Rep) for YLD confirm this fact, as Rep for YLD in 2019 is much lower than for YLD in 2020 (Supplementary Table S5). Barke, as a control, confirms that as well, as it had a yield of 127.4 dt/ha in 2019 and 60.7 dt/ha in 2020. The latter amount is consistent with the average yield of 59.5 dt/ha for Barke in a previous study in Halle (Wiegmann *et al*., 2019). In this case, HEB lines also showed lower yields than Barke.

Consequently, in addition to the ANOVA, the descriptive statistics emphasize the difference between the two trial years (Supplementary Fig. S2), meaning that differences in developmental and yield-related traits between the two years can mainly be explained by different environmental conditions (Supplementary Fig. S1, Table S4, Note S1). As a consequence, both years were evaluated separately. Correlations support this decision (Supplementary Fig. S2 and S3, Note S1). High repeatabilities and heritabilities (Supplementary Table S5) indicate that the measurements were reliable (Note S1). Furthermore, separate yearly evaluation is interesting since barley *ELF3* effects have already been shown to vary depending on the environment (Herzig *et al*., 2018).

### Comparison of HIFs reveals effects between HIF sister lines

Trait performance for each HIF line and the difference between HIF sister lines carrying the wild *ELF3_Hsp_* and the elite *ELF3_Hv_* alleles, respectively, were calculated per year (Supplementary Table S7, Fig. 2). Furthermore, descriptive statistics for each HIF sister line in each year can be found in Supplementary Table S10. For the sake of completeness, we also analysed data across years in Supplementary Table S7 and Supplementary Fig. S4.

**Figure 2.**
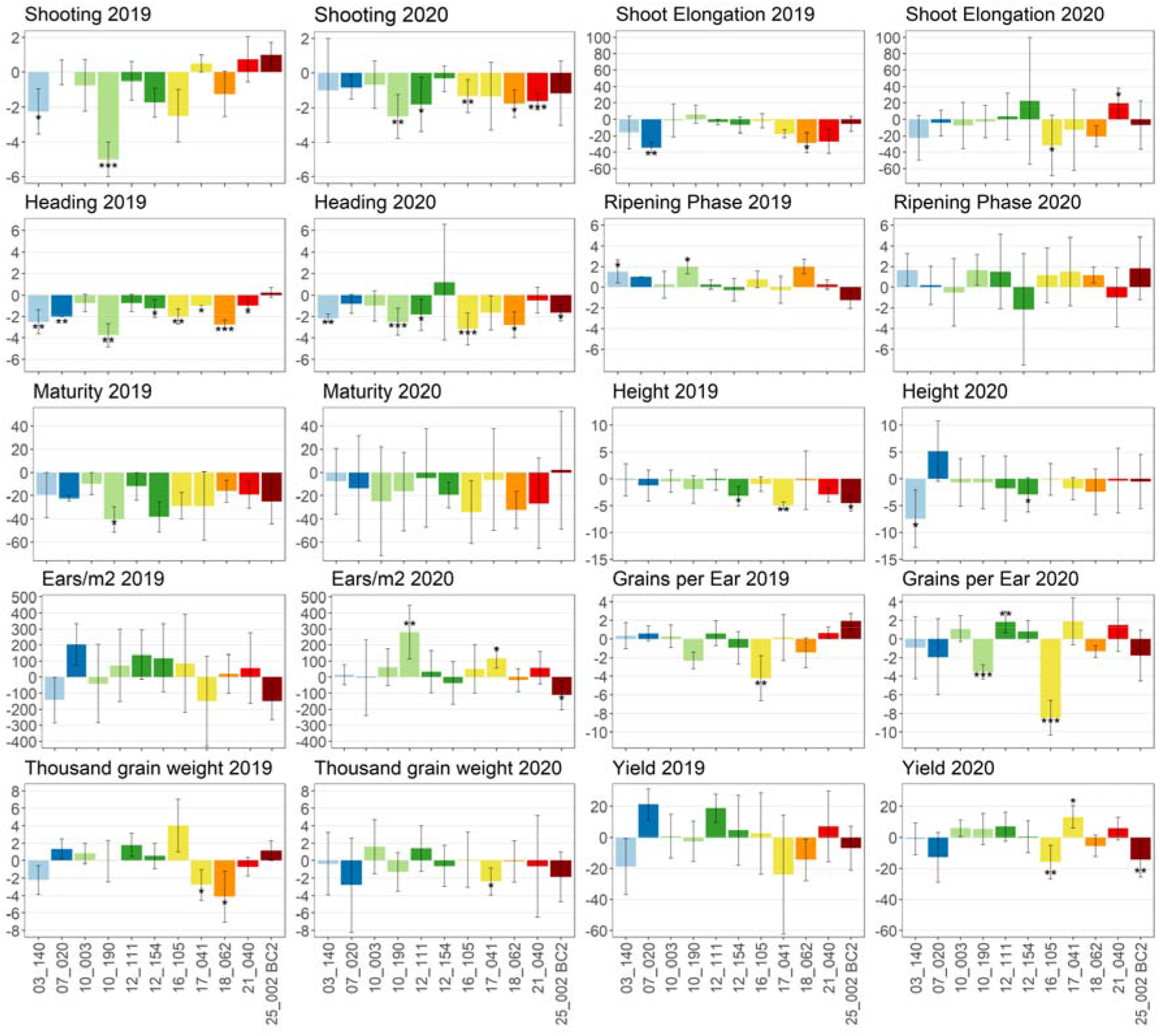
Trait differences between the two sister lines of each HIF pair (*ELF3_Hsp_* compared to *ELF3_Hv_*) per year. Lines with two identical first digits originate from the same wild donor. Trait units are given in Table 1. Asterisks indicate a significant difference between sister lines (one-way ANOVA, *p < 0.05, ** p < 0.01 and *** p < 0.001) and error bars show standard deviations. Mean differences and standard deviations are based on differences for each HIF pair per block. Columns are coloured depending on the ELF3 haplotype defined in Fig. 4C.

In general, HIF sister lines carrying an *ELF3_Hsp_* allele showed an accelerated plant development and reduced plant height in both years (Fig. 2). These findings confirm *Hsp* allele effects estimated by means of genome-wide association studies (GWAS) in previous trials (Herzig *et al*., 2018; Maurer *et al*., 2016). Also, family-specific effect variation of *ELF3_Hsp_* alleles could be seen as in Herzig *et al*. (2018). Compared to Herzig *et al*. (2018), the effects observed in the current study are more diverse and more extreme. Most likely, this is due to the relatively rough effect estimation in the GWAS model of Herzig *et al*. (2018), especially as it is based on lines with different genetic backgrounds. In the current study, the effect estimates are thought to be more reliable, as the genetic background of HIF sister lines is almost identical. In contrast to the developmental traits, the yield parameters EAR, GNE and TGW as well as YLD showed different effect directions between HIF sister lines.

Several significant effect differences were found, especially for SHO and HEA, whereof most could be confirmed in both years. The strongest and most effects were found in HIF 10_190, where SHO was up to 5.00 days earlier for the *ELF3_Hsp_* allele, for HEA up to 3.75 days earlier and for MAT up to 40.39 GDD earlier, which corresponds to 1.75 days in 2019 (Supplementary Table S7). Significant effect differences in 10_190 could also be found for the traits RIP, EAR and GNE. However, it should already be noted here, that although HIF 10_003 originates from the same wild donor (HID_102) as 10_190 and thus shares the same *ELF3_Hsp_* allele, interestingly, the strong phenotypic effects of 10_190 could not be observed in 10_003, indicating the presence of further factors determining the *ELF3_Hsp_* allele effect differences and making it necessary to carry out an inspection of the genetic background (see chapter ‘Genetic constitution of the HIF pairs beyond ELF3 impacts ELF3 effects’) and gene expression analysis (see chapter ‘Gene expression analysis reveals that phenotypic differences are not due to ELF3 transcript abundance itself’).

Yield performance of all HIFs was different between the two years. In 2020, it is striking that the absolute yield was far below average yields (Supplementary Table S7). Nevertheless, significant yield effects of *ELF3_Hsp_* allele carrying lines were found for HIFs 16_105, 17_041 and 25_002 BC2 with yield differences of up to 15.96 dt/ha in HIF 16_105. In HIF 17_041, the *ELF3_Hsp_* carrying line shows the strongest increase in yield (13.21 dt/ha in 2020). This is tremendous considering the absolute yield and the average yield for spring barley in Germany (55.6 dt/ha in 2020; Federal Ministry of Food and Agriculture, 2020). However, the correlation of earliness and yield largely depends on environmental conditions. A positive correlation of HEA and YLD in 2019 and a negative correlation in 2020 (Supplementary Fig. S3) confirm that fact and could be due to different temperature and/or precipitation resulting in a preference for later or earlier genotypes. Admittedly, a final evaluation of the influence of *ELF3* on yield would therefore require either optimal conditions without limiting factors or a series of field trials in a representative selection of environments.

Increasing yield has always been the main goal in plant breeding. Domestication and selection of crop plants improved yield but this went along with loss of genetic diversity. Wild barleys provide a huge genetic resource that can be useful to extend the elite barley breeding pool to cope with challenges set by the ongoing climate change (Ellis *et al*., 2000; Nevo, 2013; Tanksley and McCouch, 1997; Zamir, 2001). However, not only yield improving genotypes are of interest for future breeding programs, but also HIFs carrying exotic alleles for increasing biodiversity and improvement of other agronomic traits, provided that they are not associated with a yield penalty. This assumption also applies to plant height, since larger plants increase the risk of lodging and yield losses (Hedden, 2003). In this regard, depending on the environment, the *ELF3_Hsp_* carrying HIF lines 10_190 and 12_111 may be useful for breeding. The exotic alleles exhibited significantly increasing effects on EAR or GNE (Fig. 2, Supplementary Fig. S4), respectively, without simultaneous negative effects on yield or plant height. Across years, 12_111, as the only line, even had a significantly positive effect on yield (Supplementary Fig. S4). If early heading is desired, the *ELF3_Hsp_* alleles present in HIF lines 03_140 and 10_190 are interesting, since they showed early heading without negative effects on yield or plant height (Fig. 2, Supplementary Fig. S4).

However, to confirm that the found effects are actually coming from the respective *ELF3_Hsp_* alleles, an analysis of the genetic background between HIF sister lines was conducted (see chapter ‘Genetic constitution of the HIF pairs beyond ELF3 impacts ELF3 effects’).

### ELF3 effects in the context of environment

Generally, more significant trait effects were found in 2020 than in 2019 (Fig. 2). One reason could be that larger plots and more replicates (6 in 2020 vs. 4 in 2019) are necessary to observe significant differences. Another reason could be that the *ELF3_Hsp_* effect is larger under specific environmental conditions, as shown before (Herzig *et al*., 2018, Fig. 1). Herzig *et al*. (2018) reported that *ELF3_Hsp_* effects on heading were stronger in Dundee (2014 and 2015) with colder summers (up to 16 °C on average), more and equally distributed rain (>800 mm) and greater day lengths (maximum of 17.45 h) compared to Halle. In Halle the average temperature in July was up to 21 °C, 50% of the annual precipitation (514 mm) fell during July and August and maximum day length was 16.63 h (Herzig *et al*., 2018). In the present study, 2020 is characterised by a warmer vegetation period (on average 13.4 °C compared to 12.9 °C in 2019) except for the last month (on average 19.4 °C compared to 21.3 °C in 2019) with daily average temperatures of up to 23.8 °C (compared with up to 29 °C in 2019) and rain mainly at the end of the vegetation period instead of equally distributed rain as in 2019 (127 mm in both years during the vegetation period). Plants needed on average more than 11 days more to reach SHO and HEA in 2019 (Supplementary Table S8). Also, in 2020, days were longer than in 2019 for the major part of the growing season and the photoperiod (the absolute amount of day light over the whole vegetation period, CDL) was higher compared to 2019 (Supplementary Table S4). In 2019, plants on average needed more than 100 hours more of photoperiod to reach SHO and HEA (Supplementary Table S8).

As part of the circadian clock, controlling plant development based on day length and ambient temperature signals (Bendix *et al*., 2015; Calixto *et al*., 2015; Harmer, 2009; Nusinow *et al*., 2011; Wijnen and Young, 2006), *ELF3* very likely plays a role in adaptation to environmental changes in barley. In Arabidopsis, the circadian clock is a major regulator of the response to abiotic stress (reviewed in Habte *et al*. (2014)). *ELF3*, as a part of the circadian clock, might influence this as well in barley, as shown in Saade *et al*. (2016), where *ELF3_Hsp_* effects were increased under salinity stress (for HEA, TGW and HEI). *AtELF3* also controls growth in response to ambient temperature and photoperiod (Anwer *et al*., 2020; Box *et al*., 2015; Jung *et al*., 2020; Raschke *et al*., 2015; Thines and Harmon, 2010; Zhu *et al*., 2022). It was suggested to support crop improvement under higher temperature (Zhu *et al*., 2022). For barley, Ejaz and von Korff (2017) could show that a non-functional *elf3* leads to earlier flowering under high ambient temperature, whereas a functional *ELF3* leads to later flowering. Also, no reduction in floret and seed number was observed under high ambient temperature for a non-functional *elf3* compared to a functional *ELF3* allele.

Hence, we conclude that the environment in 2019 led to weaker effect differences, which could be caused by temperature, precipitation and/or photoperiod effects (Supplementary Note S1). Therefore, a further experiment under controlled greenhouse conditions was conducted.

### Image-based phenotyping in controlled environment validates results from field trials

To confirm the results from the field experiments in a different but typical experimental condition, HIF pair 10_190 was selected for a greenhouse experiment (LD: 16 h light, 8 h darkness, day/night temperatures of 20 °C/18 °C) and compared to cultivar Bowman and the two *elf3* mutant lines BW289 and BW290. The latter were generated in a Bowman background, exhibiting early flowering phenotypes (Ejaz and von Korff, 2017; Faure *et al*., 2012; Zakhrabekova *et al*., 2012). HIF pair 10_190 was selected because it exhibited the strongest effects, especially for SHO and HEA in both years in the field experiments (Fig. 2). The way of phenotyping the traits heading, tiller number and plant height was slightly different compared to the field trials. Here, heading was scored when the first awns of a plant appeared, which is well comparable with HEA in the field trials, where it was scored, when the awns were visible for 50% of all plants of a plot. In the greenhouse, number of tillers was counted manually on day 64 and all tillers were included, while in the field trials, the trait EAR was counted by using a representative 50 cm frame in the centre of a plot and only tillers actually carrying ears were counted. Plant height in the greenhouse was measured continuously and was obtained by analysing images and, in the field trial, it was solely measured at the end of maturity with tillers pulled upright.

As expected, the mutants showed earlier flowering of about 24 days compared to Bowman (Fig. 3A). For the HIF pair, the line with the wild *ELF3_Hsp_* allele flowered about 18 days earlier than the line carrying the *ELF3_Hv_* allele, even outperforming the results of the field experiments and the previous QTL studies, probably due to the optimal conditions in the greenhouse (e.g. already 16 h of light at the beginning of the experiment and a constantly optimum temperature and water supply).

**Figure 3.**
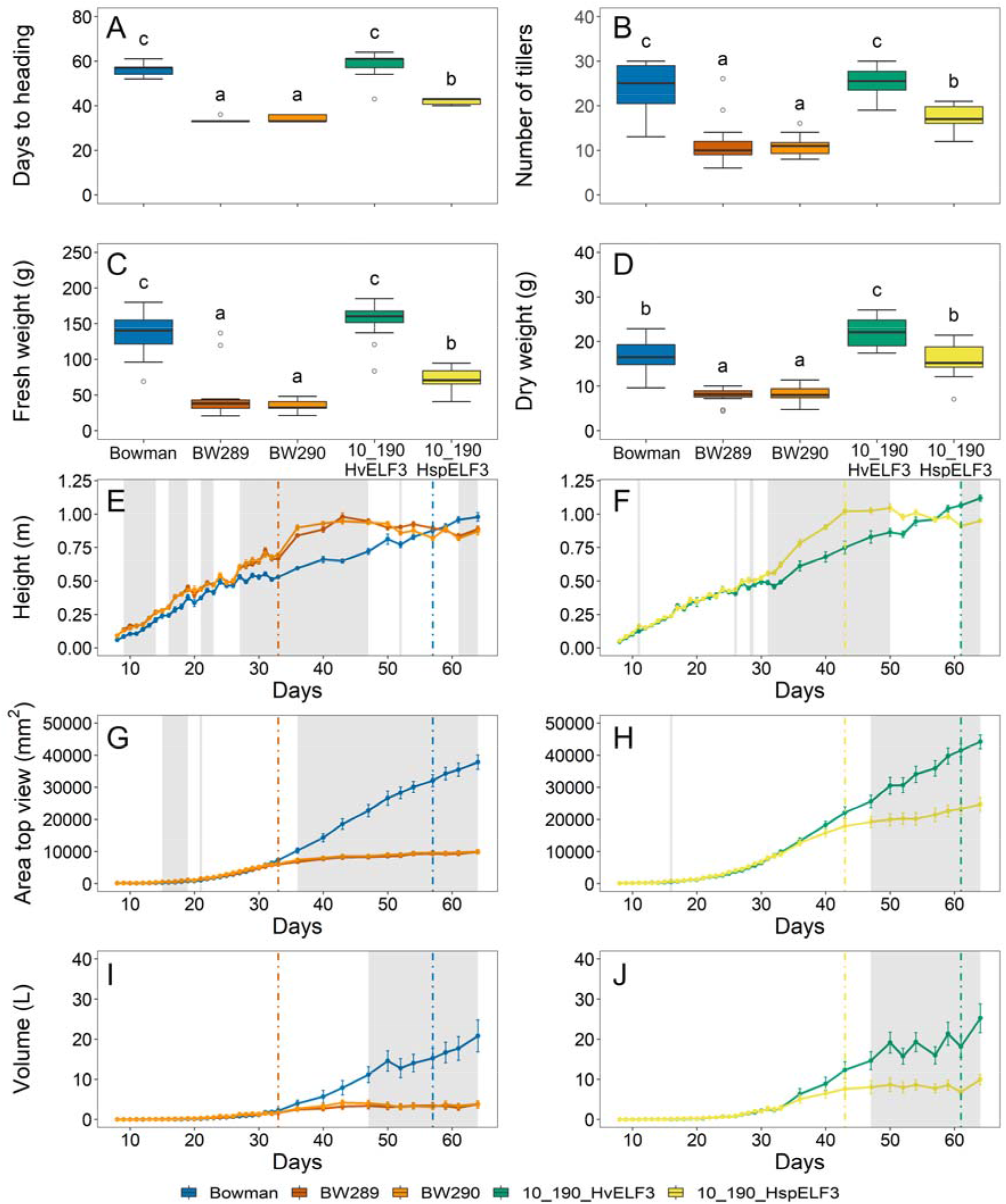
Growth and biomass parameters for cultivar Bowman, two *elf3* mutants in Bowman background, BW289 and BW290, and HIF pair 10_190. Plants were grown under standard greenhouse conditions (LD with 20/18 °C day/night temperatures). Heading (A) was scored from images when the awn tips of the first awn were visible. Number of tillers (B), fresh weight (C) and dry weight (D) were measured at the end of the experiment (day 64). Boxplots (A-D) show medians and interquartile ranges (IQR) and outliers were defined as 1.5 x IQR. Different letters above boxes indicate significant differences (one-way ANOVA, p < 0.05, Supplementary Table S11). Parameters height, area and volume (E-J) were extracted from the Integrated Analysis Platform (IAP) pipeline (Klukas *et al*., 2014). Coloured vertical lines show the mean flowering time of the respective genotype and grey shaded areas show significant differences for Bowman with both mutants (E,G,I) and between sisters lines of HIF 10_190 (F,H,J) (one-way ANOVA, p < 0.05, Supplementary Table S11). Error bars indicate standard error of mean (SEM) across ≥ 13 biological replicates.

To evaluate whether barley *ELF3* had an impact in controlling vegetative growth, the three growth parameters plant height, area and volume were measured or estimated (for volume) (Fig. 3E-J). Plant height showed an increase just before heading for both mutants and 10_190_*ELF3_Hsp_*, which could be related to the trait SHO from the field experiment where 10_190_*ELF3_Hsp_* showed early shooting (Fig. 2). Just after heading, the growth curve flattened for the mutants (day 33) and 10_190_*ELF3_Hsp_* (day 43), while growth of cultivar Bowman and 10_190_*ELF3_Hv_* continued to increase. The same trend as for plant height was visible for plant area and plant volume (Fig. 3G-J) where the growth curve flattened for the mutants and 10_190_*ELF3_Hsp_* directly after heading, whereas for Bowman and 10_190_*ELF3_Hv_* growth strongly increased at the same time. These results confirm a reduced vegetative growth rate for BW289, BW290 and the wild barley 10_190_*ELF3_Hsp_* allele. This is in accordance with the findings that BW289, BW290 and 10_190_*ELF3_Hsp_* showed fewer tillers and lower fresh and dry weight compared to cultivar Bowman and 10_190_*ELF3_Hv_* (Fig. 3B-D). This can also be explained by early heading and a shortened growth period (Fig. 3A). In the field experiment, no plant height effect was observed between sister lines of HIF 10_190 (Fig. 2). This may be explained by the fact that plant height in the field was measured at the end of maturity rather than during development. Strikingly, in the field experiment 2020, the *ELF3_Hsp_* carrying HIF 10_190 line had more ears per square meter compared to the *ELF3_Hv_* carrying line. This effect could not be validated in the greenhouse experiment. A reason could be that for the greenhouse plants, all tillers were counted without considering if tillers would develop into a spike, whereas in the field experiments only developed ears were counted. In conclusion, the greenhouse results for 10_190 were able to confirm most of the results from the field trials, in particular for heading.

### High diversity in ELF3 protein sequences

In order to understand the sequence variations of HvELF3 and to be able to better compare the Barke *ELF3_Hv_* and wild *ELF3_Hsp_* allele carrying HIF sister lines with each other, we sequenced the full-length genomic DNA of *ELF3* (Fig. 4A) from original wild barley donors and Barke. After identifying the *ELF3* coding sequence of all wild barley donors (Supplementary Table S12), the ELF3 protein sequences were determined (Supplementary Table S13) and 19 different protein types/proteoforms could be distinguished (Supplementary Table S14), whereof 9 were present in the field trials (Fig. 4C) due to the above mentioned selection criteria for HIF lines. Comparing the variation found in the wild donors of HEB-25 with already described variation for *HvELF3* in the literature revealed 23 novel mutations, whereof 13 were novel nonsynonymous SNPs (Supplementary Table S15). Furthermore, 19 *ELF3* alleles (out of 21 different *ELF3* haplotypes in donor lines of HEB-25) could be determined as novel alleles (Supplementary Table S15). Amino-terminal (N), middle (M) and carboxyl-terminal (C) regions of the HvELF3 protein were identified based on the comparison with AtELF3 (Fig. 4B), where these regions were shown to interact with different proteins (Herrero *et al*., 2012; Liu *et al*., 2001; Nieto *et al*., 2015; Yu *et al*., 2008).

**Figure 4.**
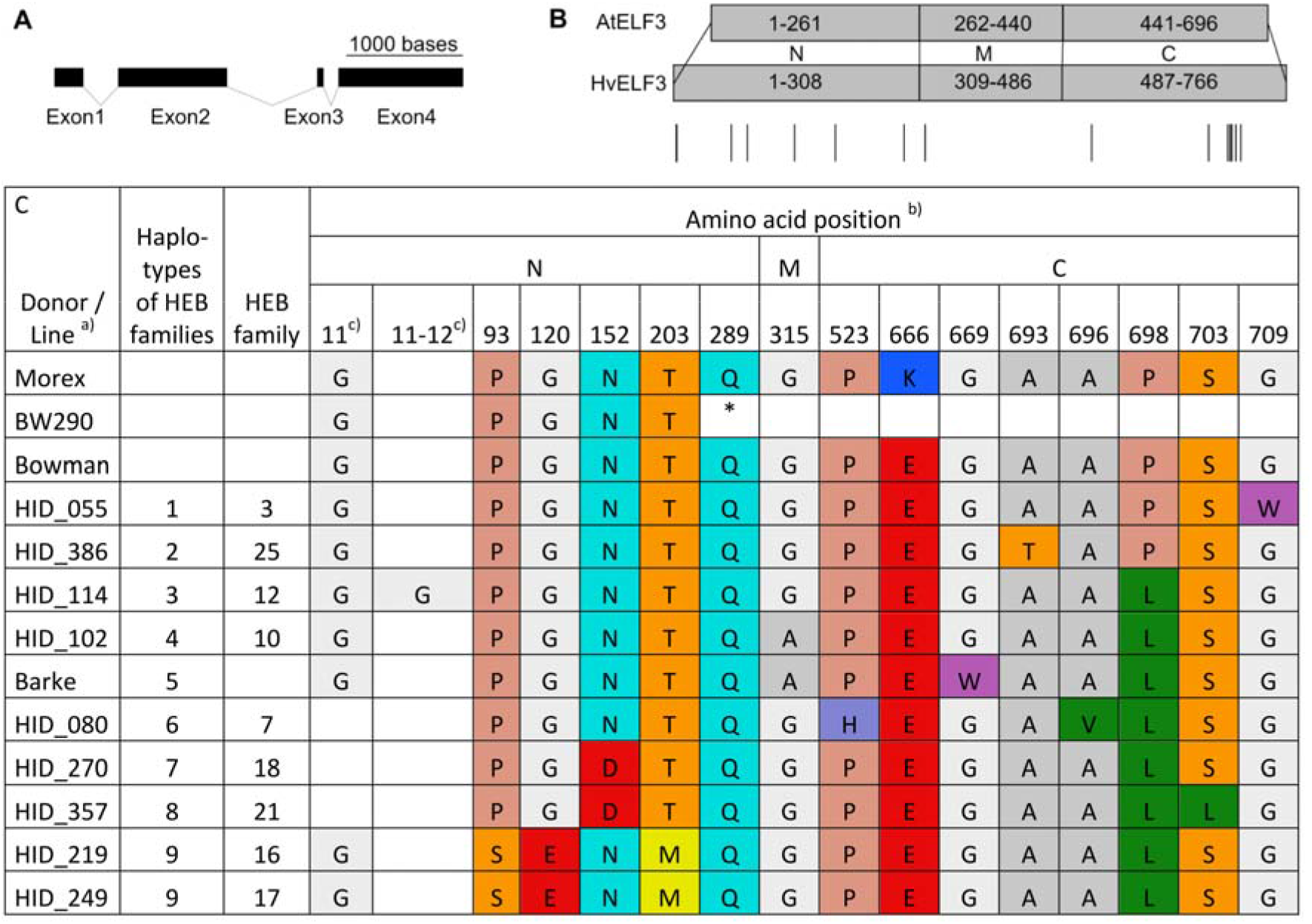
ELF3 protein structure and sequence polymorphisms. **(A)**Structure of the *HvELF3* gene in barley (Barke). Exons are shown as black rectangles and introns as connecting lines. **(B)**Domain mapping and their sequence annotation between the Arabidopsis (Col-0) ELF3 protein (AtELF3) and Barke/Morex ELF3 protein (HvELF3). Numbers indicate amino acid positions of amino-terminal (N), middle (M) and carboxyl-terminal (C) protein domains. Amino acids 696 and 766 are the STOP codons for AtELF3 and HvELF3, respectively. Lines beneath HvELF3 mark sites of amino acid substitutions and insertion or deletion between HIFs used in field trials (as indicated in C). **(C)**ELF3 protein sequence polymorphisms of all alleles present in the field trials, Morex, Bowman and BW290. Only the amino acid positions with variation between the families are shown. One-letter amino acid abbreviations (JCBN, 1984) were used and the asterisk shows a stop codon. a) HID = ‘hordeum identity’; name of the donor accession. b) N, M and C regions of barley ELF3 were obtained by alignment of the Barke/Morex sequence with the Arabidopsis sequence and Barke/Morex/Bowman sequences were used as references for the amino acid positions. c) Between position 11 and 12 some lines have an insertion and at position 11 some lines have a deletion of one amino acid, compared to the Barke amino acid sequence.

In Arabidopsis, the N region is required to interact with PHYTOCHROME B (PHYB) and CONSTITUTIVE PHOTOMORPHOGENIC 1 (COP1) (Liu *et al*., 2001; Yu *et al*., 2008), the M region with ELF4 and GIGANTEA (GI) (Herrero *et al*., 2012; Yu *et al*., 2008) and the C region with PHYTOCHROME-INTERACTING FACTOR 4 (PIF4) (Nieto *et al*., 2015). Huang *et al*. (2017) could already show that ELF3 in *Brachypodium distachyon*, a grass which is closely related to barley, interacts with almost the same set of proteins *in vivo*. While the mutation in BW289 (*eam8.k*) contains two deletions, one inversion and two small insertions (Zakhrabekova *et al*., 2012, data not shown), BW290 (*eam8.w*) has a C-to-T point mutation, resulting in a premature stop codon (Fig. 4C), leading to truncated proteins in both mutants (Faure *et al*., 2012). Since the M and C regions are absent in BW290 and since this line is flowering early (Ejaz and von Korff, 2017; Faure *et al*., 2012; Zakhrabekova *et al*., 2012), also naturally occurring mutations in these regions may influence the role of barley ELF3. Also, for the wild barley donors, most amino acid differences were observed in the N and C region. Amino acid variation at positions 315, 669 and 698 were also described in Casas *et al*. (2021) for the two cultivars Beka and Logan and suggested to be associated with differences in flowering time. Apart from that, phenotypic differences are likely to be sought also on the *cis*-regulatory level.

A summary of all ELF3 protein polymorphisms present in the field trials and their respective phenotypic effects in both years (as in Fig. 2) can be found in Supplementary Table S16. Particularly interesting is that the donors of family 16 and 17 have exactly the same protein sequence, especially when considering the different effects on yield (Fig. 2), again indicating the presence of further factors in the remaining genome determining the *ELF3_Hsp_* allele effects (see chapter ‘Genetic constitution of the HIF pairs beyond *ELF3* impacts *ELF3* effects’; especially for family 16 and 17). Moreover, the exotic ELF3 in family 10 (HID_102) only differs in one amino acid from the cultivated ELF3 of Barke. This amino acid is located at position 669 in the C-terminal region of the ELF3 protein (Fig. 4C). In Arabidopsis, the C-terminal region of ELF3 binds the PIF4 basic helix–loop–helix (bHLH) domain which subsequently prevents PIF4 from activating its transcriptional targets (Nieto *et al*., 2015). The *PIF4* gene in Arabidopsis controls thermomorphogenesis (Koini *et al*., 2009; Quint *et al*., 2016), which refers to morphological changes dependent on the ambient temperature. It regulates auxin biosynthesis, thermosensory growth, adaptations and reproductive transition (Franklin *et al*., 2011; Gangappa *et al*., 2017; Koini *et al*., 2009; Kumar *et al*., 2012). Previous studies have shown that variation in *PIF4* expression and elongation growth can be explained by genetic variation in *AtELF3* (Box *et al*., 2015; Raschke *et al*., 2015). The mutation at position 669 has already been described in the literature for other accessions (Supplementary Table S15) and has been suggested to be associated with flowering time (Casas *et al*., 2021). At this point it should also be noted, that Bowman showed the same phenotype as 10_190_*ELF3_Hv_* in the greenhouse experiment (Fig. 3) although it shares the W669G substitution as 10_190_*ELF3_Hsp_* (Fig. 4C). This and the fact that all other HIFs share this substitution (10_003 harbours even the exact same *ELF3_Hsp_* without showing strong phenotypic effect differences) strengthens the assumption that there are further factors determining the *ELF3_Hsp_* allele effect differences, like structural variation on the protein level or in the remaining genome as already suggested before. Therefore, we compared the genetic background of all HIFs and conducted gene expression analyses for *ELF3* and downstream genes in HIF 10_003 and 10_190, and performed a sequence/structure-based analysis of the ELF3_Hv_ and ELF3_Hsp_ proteins from family 10 (next chapters).

### Genetic constitution of the HIF pairs beyond ELF3 impacts ELF3 effects

To find out whether the found effects between HIF sister lines were indeed due to *ELF3_Hsp_* alleles, or may have been influenced by differences in the remaining genome, an inspection of the genetic background in HIF sister lines was carried out using the data from the 50k iSelect SNP chip (Supplementary Table S3). The genotyping results confirmed the status of the fixed homozygous *ELF3* alleles in all HIF sister lines. For the additional seven main flowering time loci found in the previous HEB-25 QTL studies (Maurer *et al*., 2015), it was possible to verify that HIF sister lines exhibited the same fixed homozygous alleles (Supplementary Table S17). Initially, we aimed for HIF pairs that would only segregate at the *ELF3* locus, but additional segregating loci between the HIF sister lines were obtained (Supplementary Fig. S5, Supplementary Table S18). Genes in these regions could possibly interfere with and have an influence on the studied traits and mask the *ELF3_Hsp_* effect. In this context, the yield-related effects in HIF sister lines of 16_105 and 17_041, which seemed interesting due to the presence of the same *ELF3* allele, can now be explained by the presence of different brittle rachis (*btr1/btr2)* alleles (physical position of the region: chromosome 3H: 38,758,057-39,626,379 bp (refseq2), closest SNP: SCRI_RS_146425, Supplementary Table S3), which affect the shattering of the ear at maturity. Thus, the observed significant yield effects are due to a differing number of harvested grains per ear and rather have to be attributed to a brittle rachis phenotype (Pourkheirandish et al., 2015) than to the *ELF3* difference. Apart from these two lines (16_105_959 (*ELF3_Hsp_*) and 17_041_1000 (*ELF3_Hv_*)), all other HIF lines carry cultivated alleles at *btr1*/*btr2* (Supplementary Table S3).

However, for a selection of genes that are already known to control flowering time or to interact with AtELF3, for example LUX and PIF4 (Nieto *et al*., 2015; Nusinow *et al*., 2011), most of the studied HIF pairs already share the same homozygous alleles (Supplementary Table S19). Nevertheless, there could be genes that are still unknown to be involved in the flowering time control pathway and the circadian clock. Of course, as few as possible additionally segregating loci are desirable. In this context, HIF pairs derived from HEB-25 lines 10_003, 10_190, 12_111 and 21_040 with only a few additional segregating regions are especially interesting (<1% of the whole genome, Supplementary Table S18). Together with the results from the field trials (Fig. 2) and the greenhouse experiment (Fig. 3), the low percentage of segregation (Supplementary Table S18 and S20) makes HIF 10_190 especially interesting.

HIFs 10_003 and 10_190 originate from the same exotic donor just like HIFs 12_111 and 12_154. Comparing these HIFs among each other regarding their genomic background, revealed contrasting alleles at five further flowering time loci for HIFs from family 10 (*PPD-H1*, *CEN*, *QFt.HEB25-4a*, *VRN-H1* and *VRN-H3/FT1* (Maurer *et al*., 2015), Supplementary Table S17) and at two further flowering time loci for HIFs from family 12 (*PPD-H1* and *VRN-H1* (Maurer *et al*., 2015), Supplementary Table S17). Also for other genes that are known to be involved in controlling flowering time, contrasting alleles were found for two *GI* related genes, *LUX*, *ELF4* and *PIF4* for HIFs in family 10 and for one *GI* related gene, *LUX*, *CO2*, *ELF4*, *PPD-H2*, *PIF4* in family 12 (Supplementary Table S19).

In the case of the HIFs in family 10, *PPD-H1* and *PIF4* are of special interest as W669G lies within the potential PIF4-interacting C region of ELF3 (Fig. 4C). In case of *PPD-H1*, the wild allele has shown the strongest influence on flowering time and also on other traits (Herzig *et al*., 2018; Maurer *et al*., 2016). Here, the *ELF3_Hsp_* effect might be increased in presence of a homozygous wild *PPD-H1* allele, suggesting an interaction of these two. A previous study has already shown increased expression of *PPD-H1* in *elf3* mutants and effects on flowering time in *elf3* mutants by variation at *PPD-H1* under LD (Faure *et al*., 2012). To verify the differences in gene expression between HIF sister lines, we measured diurnal gene expression of *ELF3* and some of its downstream genes, including *PPD-H1*.

### Gene expression analysis reveals that phenotypic differences are not due to ELF3 transcript abundance itself

To test whether *ELF3* expression is present and varies between HIF sister lines, as well as to further understand the phenotypic differences between Barke *ELF3_Hv_* and wild *ELF3_Hsp_* allele carrying HIF sister lines originating from the same wild donor, we analyzed diurnal expression of *ELF3* itself as well as its downstream genes *HvCO1*, *HvGI*, *PPD-H1*, *VRN-H3*/*HvFT1* and *VRN-H1*. Plants were grown under 16/8 h light/dark cycles with day/night temperatures of 20 °C/18 °C, and leaf samples were collected every 4 h over a 24 h period at day 17. The two HIF pairs 10_003 and 10_190 were selected due to the strong effect differences between HIF sister lines in 10_190 and the non-existent effects in 10_003 on the contrary, for which only assumptions could be made until now. Bowman and the *elf3* mutant BW290 were chosen for comparison reasons.

Bowman displayed generally higher *HvELF3* transcript abundance compared to the mutant BW290 in diurnal conditions except ZT20 (Fig. 5A), similar to previous observations under short days (SD) (Faure *et al*., 2012). However, this contradicts to another previous report using a different *elf3* mutant BW289 under 12/12h light/dark cycles (Zakhrabekova *et al*., 2012). As both BW289 and BW290 lack functional HvELF3 proteins (Zakhrabekova *et al*., 2012), the discrepancy in *HvELF3* transcript abundance is not expected to influence downstream signaling pathways. In contrast, the transcript abundance of *HvELF3* was not different between *ELF3_Hv_* and *ELF3_Hsp_* in HIFs 10_003 and 10_190, except for occasionally detected increased *HvELF3* expression in *ELF3_Hsp_* lines (10_003 at ZT20 and 10_190 at ZT00/24) (Fig. 5A). These data suggest that the observed phenotypic variation between *ELF3_Hv_* and *ELF3_Hsp_* in HIF 10_190 was not or at least not mainly due to transcript abundance of *HvELF3*, which is supported by the fact that the promotors of *ELF3_Hv_* and *ELF3_Hsp_* are identical (Supplementary Table S12).

**Figure 5.**
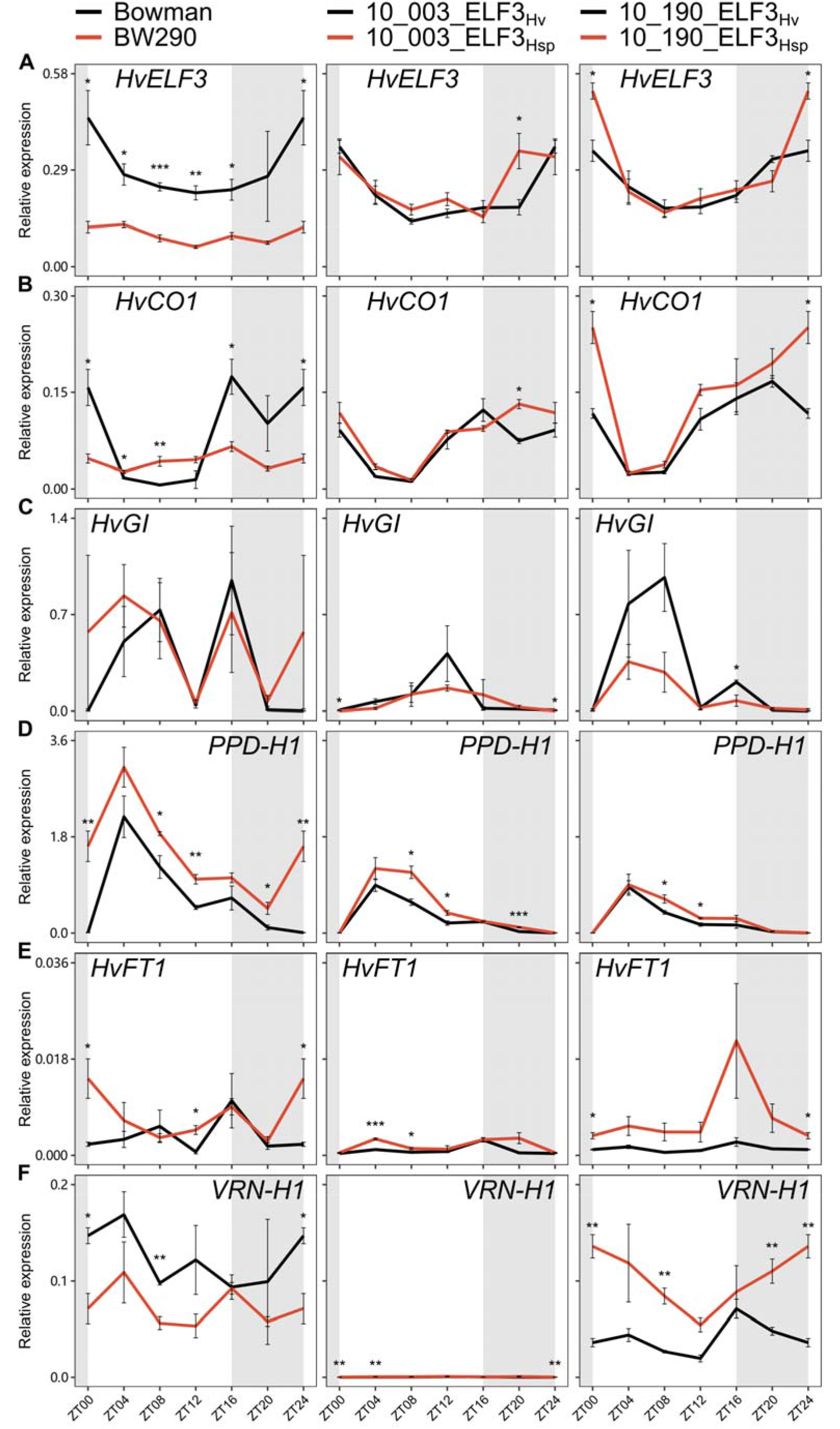
Transcript levels of flowering time genes in Bowman, BW290, HIF pairs 10_003 and 10_190. Diurnal gene expression of *HvELF3* (A), *HvCO1* (B), *HvGI (C), PPD-H1* (D), *HvFT1* (E), and *VRN-H1* (F) was measured every 4 h in plants grown under standard conditions (LD with 20/18 °C day/night temperatures). Grey-shaded areas indicate darkness. Expression levels were normalized to *HvGAPDH* and *PTF*. Error bars indicate the SEM (n=3) of three biological replicates. Asterisks above lines indicate significant differences (two-sided Student’s *t*-test, * p < 0.05, ** p < 0.01 and *** p < 0.001).

Interestingly, the rhythmic expression pattern of *HvCO1* in Bowman was not detected in BW290 (Fig. 5B). The transcript abundance of *HvCO1* in BW290 was generally higher during the light period, but lower during the dark period than in Bowman. A similar expression profile of *CO1* was found in a study with an *elf3* mutant in *Brachypodium distachyon* (Bouché *et al*., 2021), a grass closely related to barley, for which the conserved role of *ELF3* could be shown (Bouché *et al*., 2021; Huang *et al*., 2017). In that study, *CO1* expression seemed to be downregulated in the mutant during the night as in BW290. However, this strong effect of the *elf3* mutation was not observed in the HIF pairs. Both sister lines in HIFs 10_003 and 10_190 displayed rhythmic *HvCO1* expression similar to Bowman, with *ELF3_Hsp_* lines showing slightly higher expression at ZT20 in 10_003 and at ZT00/24 in 10_190. In Arabidopsis, *elf3* mutations induce *CO* expression by stabilizing GI (Yu *et al*., 2008), meanwhile the transcription of *GI* is directly repressed by the EC (Ezer *et al*., 2017). Although such ELF3-GI-CO connections are not validated in barley, *HvGI* expression was predicted to be repressed by the EC component HvLUX1 (Müller *et al*., 2020). However, almost no difference was observed in *HvGI* transcript abundance between Bowman and BW290 as well as between HIF sister lines (Fig. 5C). The observation in BW290 again contradicts the induced *HvGI* expression in a different *elf3* mutant BW289 (Zakhrabekova *et al*., 2012). Nevertheless, we could not rule out the potential effect of *ELF3* on HvGI stability, which might in turn affect *HvCO1* expression.

Consistent with previous reports (Ejaz and von Korff, 2017; Faure *et al*., 2012), the transcript abundance of *PPD-H1* increased in BW290 compared to Bowman at various time points. Such induction of *PPD-H1* expression was also observed in ELF3_Hsp_ compared to ELF3_Hv_, except for ZT00/24 in both HIFs and ZT20 in 10_190 (Fig. 5D). The fact that 10_003 and 10_190 carry differently fixed homozygous *PPD-H1_Hv_*/*PPD-H1_Hsp_*-alleles (Supplementary Table 17), responsible for later/earlier flowering, respectively (Maurer *et al*., 2016), does not seem to affect *PPD-H1* transcript abundance itself, similar as reported in an earlier study (Turner *et al*., 2005). Our data suggest that ELF3_Hsp_ in HIFs 10_003 and 10_190 induces *PPD-H1* expression regardless of its allelic differences between HIF pairs.

As expected, the increased transcript abundance of *PPD-H1* was associated with the transcriptional induction of its putative target *HvFT1* in BW290 (as in Ejaz and von Korff (2017)), 10_003_ELF3_Hsp_ and 10_190_ELF3_Hsp_ (Fig. 5E). Although *HvFT1* expression is mainly controlled by *PPD-H1* under LD, other players like *HvCO1* and/or *VRN-H1* which both upregulate *HvFT1* (Campoli *et al*., 2012a; Hemming *et al*., 2008; Turner *et al*., 2005), might be involved. The upregulation of *HvFT1* is associated with the expression of barley floral meristem identity genes *VRN-H1*, *BARLEY MADS-box 3* (*HvBM3*) and *HvBM8*, initiating inflorescence development (Digel *et al*., 2015; Ejaz and von Korff, 2017; Gol *et al*., 2021). And indeed, transcript abundance of *VRN-H1* was induced at various time points in 10_190_ELF3_Hsp_ compared to ELF3_Hv_ (Fig. 5F), consistent with its early heading phenotypes under both field and greenhouse conditions (Fig. 2 and Fig. 3). However, the *VRN-H1* transcript abundance was slightly reduced in BW290 compared to Bowman at ZT00/24 and ZT08. We would expect the altered transcript abundance of *PPD-H1* and *HvFT1* in BW290 would influence the expression of other floral regulators, such as *HvBM3* and *HvBM8*, which were more prominently correlated with *HvFT1* (Ejaz and von Korff, 2017; Gol *et al*., 2021). Although transcript abundance of *VRN-H1* was induced in 10_003_ELF3_Hsp_ compared to ELF3_Hv_, HIF 10_003 displayed overall low *VRN-H1* expression compared to Bowman, BW290, and HIF 10_190. As this HIF carries a homozygous *VRN-H1_Hsp_* from the assumed winter type wild barley donor (Supplementary Table S17), the expression of *VRN-H1* was probably repressed due to the absent vernalization. Therefore, these data can explain that the observed phenotypic differences between HIFs 10_003 and 10_190 in the field (Fig. 2) are due to different *VRN-H1* alleles.

Overall, these results confirm *ELF3* expression in the HIF lines and, as shown in previous reports with *elf3* mutants, also show differences in the expression of barley flowering time genes between HIF sister lines, although this is unlikely to be due to *ELF3* transcript abundance itself. As expected, effects of the loss-of-function *elf3* mutant on gene expression were stronger than between the natural *ELF3* variants in the HIF pairs, consistent with the respective phenotypes in the greenhouse experiment (Fig. 3). Furthermore, these results confirm the earlier assumption that the effects of *ELF3* depend on the genetic background, consistent with its upstream role in barley flowering time regulation (Boden *et al*., 2014; Faure *et al*., 2012). Therefore, comparing gene expression in the present HIF pairs is rather complicated since the HIFs originating from the same family are diverse in the remaining genome. Although it is very likely that *VRN-H1* is the reason for the huge differences between HIFs in family 10, further different homozygous alleles between HIFs 10_003 and 10_190 for *PPD-H1*, *CEN*, *QFt.HEB25-4a*, *VRN-H3/FT1* (Supplementary Table S17) and two *GI* related genes, *LUX*, *ELF4* and *PIF4* (Supplementary Table S19), several of which have been shown to interact with or be affected by ELF3, make a comprehensive comparison between these two HIFs hardly possible. This could be facilitated by the development of double HIFs (e.g. segregating at *ELF3* and *PPD-H1* or *PIF4*), with an isogenic background, in the future. Here, the concept is to detect a HEB line which is heterozygous at both loci of interest and select segregating offspring genotypes in all four possible combinations of wild and elite alleles at these two loci (i.e. *PPD-H1_Hv_*/*ELF3_Hv_*, *PPD-H1_Hv_*/*ELF3_Hsp_*, *PPD-H1_Hsp_*/*ELF3_Hv_*, *PPD-H1_Hsp_*/*ELF3_Hsp_*). Future experiments should then include phenotyping and diurnal gene expression analysis to examine whether the different combinations of wild and cultivated alleles of the respective genes show distinct effects on flowering time or other traits as well as changes in diurnal expression of themselves and downstream genes, making further statements about their interactions in a more isogenic background possible.

### W669G substitution likely affects protein structure of ELF3 and might induce disorder-driven phase separation events forming local nano-compartments

Since gene expression data suggested that the observed phenotypic variation between sister lines in HIF 10_190 was not due to *HvELF3* transcript abundance itself (Fig. 5A), it may be due to differences on the protein level, so we evaluated a potential impact of the minimal amino acid change W669G in the protein sequence between ELF3_Hv_ of Barke (ELF3_Barke_) and ELF3_Hsp_ of HID_102 (ELF3_HID102_), the exotic donor of HEB family 10. Since HIF sister lines of 10_190 are nearly isogenic (<1% of segregation, Supplementary Table S18), and it is the only HIF harbouring only one single amino acid difference between wild and Barke ELF3 (Fig. 4C) which is simultaneously showing strong phenotypic differences between HIF sister lines (Fig. 2), it is a good basis to further investigate the influence of W669G regarding its effects on the ELF3 protein structure and thereby possible influences on the phenotype. Of course, it would also be interesting to investigate the influence of other amino acid substitutions than W669G, especially when there are only differences for individual amino acids between two ELF3 proteins. One example is ELF3 in family 3 (HID_055), because it only varies from the cultivated ELF3 of Bowman at position 709 (G709W). In the context of the HIF comparison, however, the association of ELF3 variants with flowering time and crop performance would be hampered as we do not have Bowman *ELF3* alleles for comparison present in HIFs. However, this could be a future approach to create HIFs based on crossings of HID_055 and Bowman to investigate the impact of G709W on the phenotype.

In order to evaluate the potential impact of W669G on the protein level between ELF3_Barke_ and ELF3_HID102_, we performed a sequence/structure-based analysis to identify possible effects of the W669G substitution observed in HvELF3 at the protein level. Note here that computational and modelling results provide a valid and experimentally testable hypothesis and do not comprise proof. Based on InterPro (Blum *et al*., 2021), no domain is known for barley ELF3, as well as for the better annotated Arabidopsis homologue. Sensitive Markov search with HHPred (Gabler *et al*., 2020) revealed helical content with low confidence for residues 373–395. Utilizing the state-of-the-art AlphaFold2 algorithm (Jumper *et al*., 2021), the entire structure of ELF3_Barke_ and ELF3_HID102_, with the substitution was predicted (Fig. 6A,B). Interestingly, high disorder content is predicted, and, as expected, the W669G substitution is also localized in those regions (Fig. 6A,B).

**Figure 6.**
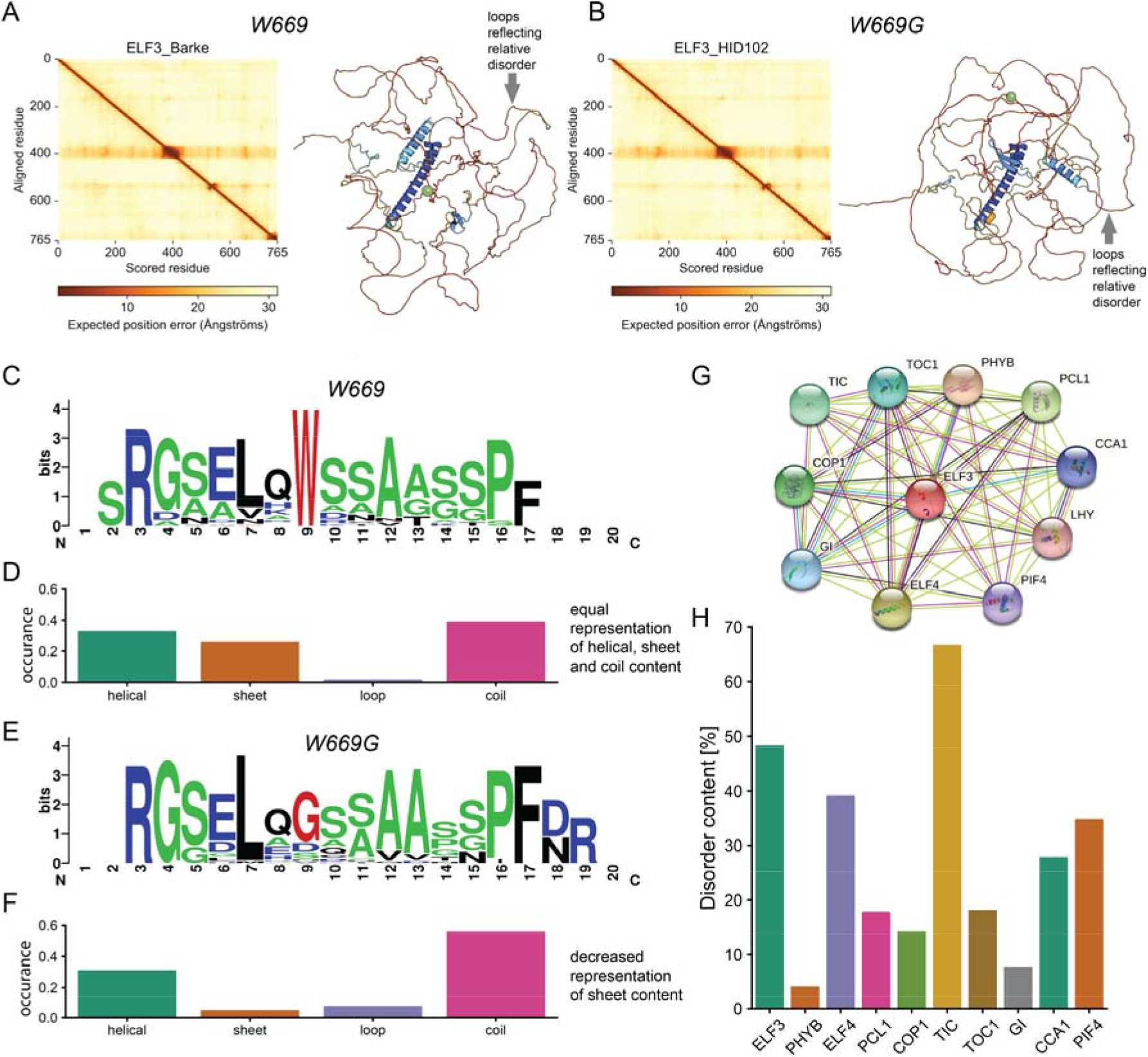
Sequence and structural analysis of the W669G substitution. (A-B) Alphafold2 prediction of the Barke and HID_102 sequence of ELF3. Models are coloured by their respective plDDT scoring, which indicates the reliability of the derived model. The Cα atom of the mutation site is highlighted as a green sphere. Left plots in (A) and (B) are standard analysis plots of Alphafold-derived models reflecting a per-residue estimate of its confidence on a scale from 0 – 100 and translated into positional ambiguity in 3D space (from 0 – 30 Å as shown in the scale bar below each plot) as described in Jumper *et al*. (2021). This measure is translated into plDDT scores and correlates with intrinsic disorder (Jumper *et al*., 2021). Red regions in the depicted Alphafold models (loops) show probable regions of high disorder content. (C-F) Local analysis of homologous structures. (C, E) Weblogo of the identified homologues for structures for Barke (W669) and substituted HID_102 (W669G) sequence. The site of the amino acid substitution at position 9 is highlighted in red, residues, prone to be in disordered regions, are highlighted in green, and charged residues in blue. (D, F) Statistical occurrence of secondary structure element in the identified structures. (G-H) Interaction analysis and disorder prediction for ELF3 and interaction partners. (G) STRING-network for ELF3 from Arabidopsis. (H) Disorder content of the barley proteins interacting with ELF3.

We next asked if the local structural preferences of this substitution could be altered. To answer this, we selected the subsequence 661-680 and structural homologues were identified by BLASTp. Identified results were filtered, using a threshold of a minimum of 5 resolved residues in the determined structure and annotated secondary structure was retrieved. In total, 52 and 34 structures were identified for the Barke and HID_102 sequence regions, respectively (Supplementary Table S21). Notably, the identified structural homologues for the Barke sequence have a conserved Tryptophan at position 9, whereas HID_102 showed lower conservation of the substituted Glycine (Fig. 6C-F). Analysing the secondary structure content of identified homologues revealed deviations in local folding preference (Fig. 6D,F, Supplementary Table S21). For Barke, the region folds in α-helical, β-sheet and coil conformations with equal occurrence. For the variant W669G in HID_102, a dramatic decrease in β-sheet occurrence is derived (Fig. 6F). Glycine, besides Proline, is a known β-sheet breaker (Minor and Kim, 1994; Smith *et al*., 1994), corroborating this observation with possible effects on higher-order hydrogen bonding patterns.

ELF3 contains high content of unstructured/disordered protein regions (Fig. 6A) and, locally, these regions can transition equally to various secondary structure elements as shown by our analysis (Fig. 6D,F). Disordered protein regions are often linked to phase separation events (Majumdar *et al*., 2019), forming local nano-compartments in the cell. Given the fact that AtELF3 itself phase-separates (Jung *et al*., 2020), this substitution could well affect the phase separation behaviour of ELF3 in barley, i.e. the switch between the active and the inactive state, as shown for Arabidopsis in the context of temperature sensitivity (Jung *et al*., 2020). The causal relationship and functional consequences of phase separation *in vivo* are very elusive (McSwiggen *et al*., 2019), so there are still many open questions regarding potential effects and functional consequences of ELF3 phase separation and nano-compartment formation in barley. Another, less obvious effect of W669G could be in the context of its local cellular interactions, and, we therefore analysed its annotated cellular community. Based on the STRING (Szklarczyk *et al*., 2021) entry of the homologous protein from Arabidopsis, AtELF3 has 10 described interaction partners identified using the default criteria (Fig. 6G). We identified the respective barley proteins using BLASTp and analysed the disorder content utilizing the MobiDB-lite algorithm (Necci *et al*., 2017) (Supplementary Table S22). The majority of the interaction partners show high disorder content as well with a mean value of 28% when considering their full sequences (Fig. 6H). Based on this, the effect of this W669G substitution might not only affect the phase separation behaviour of ELF3, but might also be involved in disorder-driven phase separation events within its cellular community.

The substitution of residue 669 from Tryptophane to Glycine might play an essential role in regulating a function in a higher-order assembly. This is because (a) the Tryptophane-containing sequence can adopt all types of secondary structure, whereas the Glycine substitution induces a reduced β-sheet content due to the sheet-breaking properties of Glycine. This might directly affect secondary structure transitions, needed in disordered regions to adapt for self-interacting (as in the case of AtELF3 (Jung *et al*., 2020)) or/and interacting with different interaction partners and thereby influencing complex composition and higher-order community 3D architecture (Kim and Han, 2018); (b) Tryptophane contains a delocalized π-electron system in its side chain, and is thereby able to form π-π, π-stacking and cation-π interaction networks. This interaction seems to play an essential role in the process of phase separation (Vernon *et al*., 2018). The analysed substitution might thereby directly affect the properties underlying nano-compartment formation and, ultimately, regulate a functional complex to perform function with distinct phenotypic consequences.

However, the fact that *Hsp* alleles of all other HIFs also share this substitution (Fig. 4) and 10_003, which harbours exactly the same *ELF3_Hsp_* allele without showing the same strong phenotypic effects, underline the dependency of the *ELF3* effect on the genetic background, shaping a complex interaction network.

## Conclusions and future prospects

In this study, we validated QTL effects from previous barley field studies that were attributed to a genomic region that included *ELF3* (Herzig *et al*., 2018; Maurer *et al*., 2015; Maurer *et al*., 2016). We determined novel exotic *ELF3* alleles and made use of nearly isogenic barley HIF pairs that segregated for the *ELF3* gene. The HIF pairs confirmed variation between *ELF3* alleles and genotype by environment interaction across the studied years. A greenhouse experiment confirmed the major results of the field trials. HIF pair 10_190 was especially promising, showing strong effects without yield losses in the field experiment and even stronger effects in the greenhouse. For natural *ELF3* variants, we found variation regarding flowering time gene expression in barley. However, the phenotypic trait differences could not be explained by differences in *ELF3* transcript abundance itself, but may be explained by the substitution of a single amino acid, which was predicted to influence ELF3 protein structure, thereby possibly affecting the properties underlying nano-compartment formation of ELF3 and potentially also affecting its interaction partners inside the cell. This emphasizes the influence of further factors impacting the phenotypic *ELF3_Hsp_* allele effects, like further variation at the protein level or in the remaining genome, e.g. alleles at other genes like *PPD-H1*, *PIF4* or *VRN-H1*, for some of which a modified expression could already be shown (Fig. 5). Due to the central role of ELF3 in the circadian clock with manifold protein interactions, in future experiments additional HIFs differing in the genomic background should be selected and characterized to shed further light on the control of plant development by interacting substitutions at critical amino acid positions of the ELF3 protein. Ultimately, this study confirmed that HIFs can be a useful tool to characterize and validate allelic effects from previous QTL studies. We have shown that the selection of HIFs with a fixed genomic background is crucial to obtain significant results. Furthermore, we propose double HIFs, simultaneously segregating at two loci, as a valuable option to investigate epistatic effects or dependencies between interacting genes. The identification of promising *ELF3* alleles for improvement of developmental and yield-related traits in barley is important for barley breeding, especially for adaptation of elite barley to climate change related stresses.

## Supplementary data

*Table S1*. IBD Genotype data from the Infinium iSelect 50k SNP chip of preselected BC_1_S_3:8_ lines with a heterozygous *ELF3* locus.

*Table S2*. Markers used for selection of HIF sister lines.

*Table S3*. IBD Genotype data from the Infinium iSelect 50k SNP chip of all HIFs used in the field trial (with a homozygous *ELF3* locus).

*Table S4*. Day lengths and weather conditions during the respective growth periods in 2019 and 2020.

*Table S5*. Repeatabilies (Rep) and heritabilites (H^2^).

*Table S6*. Primers used for PCR and sequencing of *ELF3* coding and promotor sequence and primers used for gene expression analysis.

*Table S7*. Raw data and BLUEs of all investigated traits with significant differences between HIF sister lines.

*Table S8*. Descriptive statistics of all investigated traits based on best linear unbiased estimates (BLUEs) for both years separately.

*Table S9*. ANOVA results of phenotypic data for genotype, year and genotype×year interactions.

*Table S10*. Descriptive statistics for each HIF sister line per year.

*Table S11*. ANOVA results of phenotypic data for image-based greenhouse experiment.

*Table S12*. Coding sequences of all 25 wild donors of HEB-25, Barke, Bowman and BW290 and promotor sequence of *ELF3* in HIF sister lines of 10_190.

*Table S13*. Protein sequences of all 25 wild donors of HEB-25, Barke, Bowman, BW290 and Morex.

*Table S14*. Variation in amino acids of all 25 wild donors of HEB-25, Barke, Bowman, BW290 and Morex.

*Table S15*. Variation in coding sequences of all 25 wild donors of HEB-25, Barke, Bowman and BW290 and comparison with variation already described in the literature.

*Table S16*. Summarizing table of all ELF3 protein sequence polymorphisms present in the field trials and their respective phenotypic effects in both years.

*Table S17*. IBD Genotype data from the Infinium iSelect 50k SNP chip of all HIFs for the eight major flowering time loci.

*Table S18*. Segregation of HIF sister lines in basepairs and in % of whole barley genome of 5.1 Gbp including the ELF3 region.

*Table S19*. IBD Genotype data from the Infinium iSelect 50k SNP chip of a selection of genes that are also important for flowering time control in barley.

*Table S20*. Genes in segregating regions of line 10_190 extracted from Barlex (IPK Gatersleben).

*Table S21*. Data from the local sequence analysis, including the identified pdb file, chain, residue range and occurrence of secondary structure elements.

*Table S22*. Accession codes for the barley homologues and the disorder content prediction.

*Fig. S1*. Weather data.

*Fig. S2*. Boxplots for all traits and both years separately.

*Fig. S3*. Correlations of traits for 2019 and 2020 separately.

*Figure S4*. Trait differences between the two sister lines of each HIF pair (*ELF3_Hsp_* compared to *ELF3_Hv_*) across years.

*Fig. S5*. Segregating regions between HIF sister lines.

*Note S1*. Repeatabilities and heritabilities. Correlations. Impacts of the environment on phenotypes.

## Acknowledgements

This work was funded by the European Social Fund (ESF) through the AGRIPOLY Graduate School *“Determinants of Plant Performance”* (project leaders: KP and MQ). We want to thank Matthias Reimers, Kathrin Denk, and Jana Trenner for their assistance in gene expression analysis. We are also grateful to Roswitha Ende, Jana Müglitz, Markus Hinz and various students for technical assistance in the field experiments. Furthermore, we want to thank TraitGenetics GmbH, Gatersleben, Germany, for genotyping HIF sister lines with KASP markers and with the barley Infinium iSelect 50k SNP chip. We also want to thank the Leibniz Institute of Plant Genetics and Crop Plant Research (IPK), Gatersleben, Germany, for the possibility to conduct an image-based phenotyping experiment and especially Ingo Mücke, Annett Busching, Gunda Wehrstedt, Marie Cheyenne Hellmann and Heiko Kriegel for their support with the experiment. PLK thanks the Federal Ministry for Education and Research (BMBF, ZIK program) (Grant nos. 03Z22HN23, 03Z22HI2 and 03COV04 to PLK), the European Regional Development Funds for Saxony-Anhalt (grant no. EFRE: ZS/2016/04/78115 to PLK), the Deutsche Forschungsgemeinschaft (DFG) (project number 391498659 and RTG 2467), and the Martin Luther University of Halle-Wittenberg.

## Author contributions

KP, MQ and AM conceived the project and planned the experiments. TZ analysed all data and performed the experiment in 2019. JK and NR performed the experiment in 2020. ZZ generated *ELF3* gene sequences, derived ELF3 protein sequences and conceived the image-based phenotyping experiment, which was conducted by AJ and TA. ZZ and SB performed gene expression experiments and SB generated *ELF3* promotor sequences of HIF 10_190. TS analysed *ELF3* gene sequences and provided additional barley genome resources. CT and PLK performed ELF3 protein sequence and structure analysis. TZ, ZZ, CT, MQ, KP and AM wrote the manuscript.

## Data availability

All relevant data are included in the supplementary files. Protein sequence/structure analysis scripts can be made available upon request.

## References

Alqudah AM, Schnurbusch T. 2017. Heading Date Is Not Flowering Time in Spring Barley. Front Plant Sci 8, 896.

Andres F, Coupland G. 2012. The genetic basis of flowering responses to seasonal cues. Nature Review Genetics 13, 627–639.

Anwer MU, Davis A, Davis SJ, Quint M. 2020. Photoperiod sensing of the circadian clock is controlled by EARLY FLOWERING 3 and GIGANTEA. Plant Journal 101, 1397–1410.

Bayer MM, Rapazote-Flores P, Ganal M, et al. 2017. Development and Evaluation of a Barley 50k iSelect SNP Array. Frontiers in Plant Science 8, 1792.

Bendix C, Marshall CM, Harmon FG. 2015. Circadian Clock Genes Universally Control Key Agricultural Traits. Molecular Plant 8, 1135–1152.

Bergelson J, Roux F. 2010. Towards identifying genes underlying ecologically relevant traits in Arabidopsis thaliana. Nature Review Genetics 11, 867–879.

Bhatla N. 2012. Exon-Intron Graphic Maker. http://wormweb.org/exonintron. Accessed November 2021.

Blum M, Chang HY, Chuguransky S, et al. 2021. The InterPro protein families and domains database: 20 years on. Nucleic Acids Research 49, D344–D354.

Blümel M, Dally N, Jung C. 2015. Flowering time regulation in crops-what did we learn from Arabidopsis? Current Opinion in Biotechnology 32, 121–129.

Boden SA, Weiss D, Ross JJ, Davies NW, Trevaskis B, Chandler PM, Swain SM. 2014. EARLY FLOWERING3 Regulates Flowering in Spring Barley by Mediating Gibberellin Production and FLOWERING LOCUS T Expression. Plant Cell 26, 1557–1569.

Bouché F, Woods DP, Linden J, Li W, Mayer KS, Amasino RM, Perilleux C. 2021. EARLY FLOWERING 3 and Photoperiod Sensing in Brachypodium distachyon. Front Plant Sci 12, 769194.

Box MS, Huang BE, Domijan M, et al. 2015. ELF3 controls thermoresponsive growth in Arabidopsis. Current Biology 25, 194–199.

Calixto CP, Waugh R, Brown JW. 2015. Evolutionary relationships among barley and Arabidopsis core circadian clock and clock-associated genes. Journal of Molecular Evolution 80, 108–119.

Cammarano D, Ronga D, Francia E, et al. 2021. Genetic and Management Effects on Barley Yield and Phenology in the Mediterranean Basin. Front Plant Sci 12, 655406.

Campoli C, Drosse B, Searle I, Coupland G, von Korff M. 2012a. Functional characterisation of HvCO1, the barley (Hordeum vulgare) flowering time ortholog of CONSTANS. Plant Journal 69, 868–880.

Campoli C, Shtaya M, Davis SJ, von Korff M. 2012b. Expression conservation within the circadian clock of a monocot: natural variation at barley Ppd-H1 affects circadian expression of flowering time genes, but not clock orthologs. BMC Plant Biology 12, 97.

Campoli C, von Korff M. 2014. Genetic Control of Reproductive Development in Temperate Cereals. Advances in Botanical Research 72, 131–158.

Casas AM, Gazulla CR, Monteagudo A, et al. 2021. Candidate genes underlying QTL for flowering time and their interactions in a wide spring barley (Hordeum vulgare L.) cross. The Crop Journal 9, 862–872.

Challinor AJ, Watson J, Lobell DB, Howden SM, Smith DR, Chhetri N. 2014. A meta-analysis of crop yield under climate change and adaptation. Nature Climate Change 4, 287–291.

Cockram J, Jones H, Leigh FJ, O’Sullivan D, Powell W, Laurie DA, Greenland AJ. 2007. Control of flowering time in temperate cereals: genes, domestication, and sustainable productivity. Journal of Experimental Botany 58, 1231–1244.

Crooks GE, Hon G, Chandonia JM, Brenner SE. 2004. WebLogo: a sequence logo generator. Genome Research 14, 1188–1190.

Deng W, Casao MC, Wang P, Sato K, Hayes PM, Finnegan EJ, Trevaskis B. 2015. Direct links between the vernalization response and other key traits of cereal crops. Nature Communications 6, 5882.

Digel B, Pankin A, von Korff M. 2015. Global Transcriptome Profiling of Developing Leaf and Shoot Apices Reveals Distinct Genetic and Environmental Control of Floral Transition and Inflorescence Development in Barley. Plant Cell 27, 2318–2334.

Dixon LE, Knox K, Kozma-Bognar L, Southern MM, Pokhilko A, Millar AJ. 2011. Temporal repression of core circadian genes is mediated through EARLY FLOWERING 3 in Arabidopsis. Current Biology 21, 120–125.

Ejaz M, von Korff M. 2017. The Genetic Control of Reproductive Development under High Ambient Temperature. Plant Physiology 173, 294–306.

Ellis RP, Forster BP, Robinson D, Handley LL, Gordon DC, Russell JR, Powell W. 2000. Wild barley: a source of genes for crop improvement in the 21st century? Journal of Experimental Botany 51, 9–17.

Ezer D, Jung JH, Lan H, et al. 2017. The evening complex coordinates environmental and endogenous signals in Arabidopsis. Nat Plants 3, 17087.

FAOSTAT. 2009. High Level Expert Forum - How to feed the world 2050.

Faure S, Turner AS, Gruszka D, Christodoulou V, Davis SJ, von Korff M, Laurie DA. 2012. Mutation at the circadian clock gene EARLY MATURITY 8 adapts domesticated barley (Hordeum vulgare) to short growing seasons. Proceedings of the National Academy of Sciences of the United States of America 109, 8328–8333.

Federal Ministry of Food and Agriculture. 2020. Erntebericht 2020 - Mengen und Preise.

Fernandez-Calleja M, Casas AM, Igartua E. 2021. Major flowering time genes of barley: allelic diversity, effects, and comparison with wheat. Theoretical and Applied Genetics 134, 1867–1897.

Francia E, Tondelli A, Rizza F, et al. 2011. Determinants of barley grain yield in a wide range of Mediterranean environments. Field Crops Research 120, 169–178.

Franklin KA, Lee SH, Patel D, et al. 2011. Phytochrome-interacting factor 4 (PIF4) regulates auxin biosynthesis at high temperature. Proceedings of the National Academy of Sciences of the United States of America 108, 20231–20235.

Gabler F, Nam SZ, Till S, Mirdita M, Steinegger M, Soding J, Lupas AN, Alva V. 2020. Protein Sequence Analysis Using the MPI Bioinformatics Toolkit. Current Protocols in Bioinformatics 72, e108.

Gangappa SN, Berriri S, Kumar SV. 2017. PIF4 Coordinates Thermosensory Growth and Immunity in Arabidopsis. Current Biology 27, 243–249.

Gasteiger E, Gattiker A, Hoogland C, Ivanyi I, Appel RD, Bairoch A. 2003. ExPASy: The proteomics server for in-depth protein knowledge and analysis. Nucleic Acids Research 31, 3784–3788.

Gol L, Haraldsson EB, von Korff M. 2021. Ppd-H1 integrates drought stress signals to control spike development and flowering time in barley. Journal of Experimental Botany 72, 122–136.

Habte E, Muller LM, Shtaya M, Davis SJ, von Korff M. 2014. Osmotic stress at the barley root affects expression of circadian clock genes in the shoot. Plant Cell Environment 37, 1321–1327.

Harmer SL. 2009. The circadian system in higher plants. Annual Review of Plant Biology 60, 357–377.

Hedden P. 2003. The genes of the Green Revolution. Trends in Genetics 19, 5–9.

Hemming MN, Peacock WJ, Dennis ES, Trevaskis B. 2008. Low-temperature and daylength cues are integrated to regulate FLOWERING LOCUS T in barley. Plant Physiology 147, 355–366.

Herrero E, Kolmos E, Bujdoso N, et al. 2012. EARLY FLOWERING4 recruitment of EARLY FLOWERING3 in the nucleus sustains the Arabidopsis circadian clock. Plant Cell 24, 428–443.

Herzig P, Backhaus A, Seiffert U, von Wiren N, Pillen K, Maurer A. 2019. Genetic dissection of grain elements predicted by hyperspectral imaging associated with yield-related traits in a wild barley NAM population. Plant Science 285, 151–164.

Herzig P, Maurer A, Draba V, et al. 2018. Contrasting genetic regulation of plant development in wild barley grown in two European environments revealed by nested association mapping. Journal of Experimental Botany 69, 1517–1531.

Hicks KA, Albertson TM, Wagner DR. 2001. EARLY FLOWERING3 encodes a novel protein that regulates circadian clock function and flowering in Arabidopsis. Plant Cell 13, 1281–1292.

Huang H, Alvarez S, Bindbeutel R, et al. 2016. Identification of Evening Complex Associated Proteins in Arabidopsis by Affinity Purification and Mass Spectrometry. Molecular & Cellular Proteomics 15, 201–217.

Huang H, Gehan MA, Huss SE, et al. 2017. Cross-species complementation reveals conserved functions for EARLY FLOWERING 3 between monocots and dicots. Plant Direct 1, e00018.

Huang H, Nusinow DA. 2016. Into the Evening: Complex Interactions in the Arabidopsis Circadian Clock. Trends in Genetics 32, 674–686.

Jayakodi M, Padmarasu S, Haberer G, et al. 2020. The barley pan-genome reveals the hidden legacy of mutation breeding. Nature 588, 284–289.

JCBN I-IJCoBN. 1984. Nomenclature and symbolism for amino acids and peptides. Recommendations 1983. European Journal of Biochemistry 138, 9–37.

Jumper J, Evans R, Pritzel A, et al. 2021. Highly accurate protein structure prediction with AlphaFold. Nature 596, 583–589.

Jung JH, Barbosa AD, Hutin S, et al. 2020. A prion-like domain in ELF3 functions as a thermosensor in Arabidopsis. Nature 585, 256–260.

Katoh K, Rozewicki J, Yamada KD. 2019. MAFFT online service: multiple sequence alignment, interactive sequence choice and visualization. Briefings in Bioinformatics 20, 1160–1166.

Kazan K, Lyons R. 2016. The link between flowering time and stress tolerance. J Exp Bot 67, 47–60.

Kim DH, Han KH. 2018. Transient Secondary Structures as General Target-Binding Motifs in Intrinsically Disordered Proteins. International Journal of Molecular Sciences 19.

Klukas C, Chen D, Pape JM. 2014. Integrated Analysis Platform: An Open-Source Information System for High-Throughput Plant Phenotyping. Plant Physiology 165, 506–518.

Koini MA, Alvey L, Allen T, Tilley CA, Harberd NP, Whitelam GC, Franklin KA. 2009. High temperature-mediated adaptations in plant architecture require the bHLH transcription factor PIF4. Current Biology 19, 408–413.

Kumar SV, Lucyshyn D, Jaeger KE, Alos E, Alvey E, Harberd NP, Wigge PA. 2012. Transcription factor PIF4 controls the thermosensory activation of flowering. Nature 484, 242–245.

Kuraku S, Zmasek CM, Nishimura O, Katoh K. 2013. aLeaves facilitates on-demand exploration of metazoan gene family trees on MAFFT sequence alignment server with enhanced interactivity. Nucleic Acids Research 41, W22–28.

Liu XL, Covington MF, Fankhauser C, Chory J, Wagner DR. 2001. ELF3 encodes a circadian clock-regulated nuclear protein that functions in an Arabidopsis PHYB signal transduction pathway. Plant Cell 13, 1293–1304.

Majumdar A, Dogra P, Maity S, Mukhopadhyay S. 2019. Liquid-Liquid Phase Separation Is Driven by Large-Scale Conformational Unwinding and Fluctuations of Intrinsically Disordered Protein Molecules. Journal of Physical Chemistry Letters 10, 3929–3936.

Maurer A, Draba V, Jiang Y, Schnaithmann F, Sharma R, Schumann E, Kilian B, Reif JC, Pillen K. 2015. Modelling the genetic architecture of flowering time control in barley through nested association mapping. BMC Genomics 16, 290.

Maurer A, Draba V, Pillen K. 2016. Genomic dissection of plant development and its impact on thousand grain weight in barley through nested association mapping. Journal of Experimental Botany 67, 2507–2518.

Maurer A, Pillen K. 2019. 50k Illumina Infinium iSelect SNP Array data for the wild barley NAM population HEB-25. doi: 10.5447/ipk/2019/20.

Maurer A, Sannemann W, Leon J, Pillen K. 2017. Estimating parent-specific QTL effects through cumulating linked identity-by-state SNP effects in multiparental populations. Heredity 118, 477–485.

McMaster GS, Wilhelm WW. 1997. Growing degree-days: one equation, two interpretations Agricultural and Forest Meteorology 87, 291–300.

McSwiggen DT, Mir M, Darzacq X, Tjian R. 2019. Evaluating phase separation in live cells: diagnosis, caveats, and functional consequences. Genes Dev 33, 1619–1634.

Minor DL, Jr., Kim PS. 1994. Measurement of the beta-sheet-forming propensities of amino acids. Nature 367, 660–663.

Monat C, Padmarasu S, Lux T, et al. 2019. TRITEX: chromosome-scale sequence assembly of Triticeae genomes with open-source tools. Genome Biology 20, 284.

Müller LM, Mombaerts L, Pankin A, Davis SJ, Webb AAR, Goncalves J, von Korff M. 2020. Differential Effects of Day/Night Cues and the Circadian Clock on the Barley Transcriptome. Plant Physiology 183, 765–779.

Necci M, Piovesan D, Dosztanyi Z, Tosatto SCE. 2017. MobiDB-lite: fast and highly specific consensus prediction of intrinsic disorder in proteins. Bioinformatics 33, 1402–1404.

Nevo E. 2013. Evolution of Wild Barley and Barley Improvement. Dordrecht: Springer Netherlands, 1–23.

Nieto C, Lopez-Salmeron V, Daviere JM, Prat S. 2015. ELF3-PIF4 interaction regulates plant growth independently of the Evening Complex. Current Biology 25, 187–193.

Nusinow DA, Helfer A, Hamilton EE, King JJ, Imaizumi T, Schultz TF, Farre EM, Kay SA. 2011. The ELF4-ELF3-LUX complex links the circadian clock to diurnal control of hypocotyl growth. Nature 475, 398–402.

Pourkheirandish M, Hensel G, Kilian B, et al. 2015. Evolution of the Grain Dispersal System in Barley. Cell, 527–539.

Quint M, Delker C, Franklin KA, Wigge PA, Halliday KJ, van Zanten M. 2016. Molecular and genetic control of plant thermomorphogenesis. Nature Plants 2, 15190.

Raschke A, Ibanez C, Ullrich KK, et al. 2015. Natural variants of ELF3 affect thermomorphogenesis by transcriptionally modulating PIF4-dependent auxin response genes. BMC Plant Biology 15, 197.

Saade S, Maurer A, Shahid M, Oakey H, Schmockel SM, Negrao S, Pillen K, Tester M. 2016. Yield-related salinity tolerance traits identified in a nested association mapping (NAM) population of wild barley. Scientific Reports 6, 32586.

Semagn K, Babu R, Hearne S, Olsen M. 2014. Single nucleotide polymorphism genotyping using Kompetitive Allele Specific PCR (KASP): overview of the technology and its application in crop improvement. Molecular Breeding 33.

Smith CK, Withka JM, Regan L. 1994. A thermodynamic scale for the beta-sheet forming tendencies of the amino acids. Biochemistry 33, 5510–5517.

Szklarczyk D, Gable AL, Nastou KC, et al. 2021. The STRING database in 2021: customizable protein-protein networks, and functional characterization of user-uploaded gene/measurement sets. Nucleic Acids Research 49, D605–D612.

Tanksley SD, McCouch SR. 1997. Seed banks and molecular maps: unlocking genetic potential from the wild. Science 277, 1063–1066.

The Arabidopsis Information Resource (TAIR). https://www.arabidopsis.org/servlets/TairObject?type=locus&name=at2g25930. Accessed November 29, 2021.

Thines B, Harmon FG. 2010. Ambient temperature response establishes ELF3 as a required component of the core Arabidopsis circadian clock. Proceedings of the National Academy of Sciences of the United States of America 107, 3257–3262.

Tuinstra MR, Ejeta G, Goldsbrough PB. 1997. Heterogeneous inbred family (HIF) analysis: a method for developing near-isogenic lines that differ at quantitative trait loci. Theoretical and Applied Genetics 95, 1005–1011.

Turner A, Beales J, Faure S, Dunford RP, Laurie DA. 2005. The pseudo-response regulator Ppd-H1 provides adaptation to photoperiod in barley. Science 310, 1031–1034.

Vernon RM, Chong PA, Tsang B, Kim TH, Bah A, Farber P, Lin H, Forman-Kay JD. 2018. Pi-Pi contacts are an overlooked protein feature relevant to phase separation. Elife 7.

von Zitzewitz J, Szucs P, Dubcovsky J, et al. 2005. Molecular and structural characterization of barley vernalization genes. Plant Molecular Biology 59, 449–467.

Wiegmann M, Maurer A, Pham A, et al. 2019. Barley yield formation under abiotic stress depends on the interplay between flowering time genes and environmental cues. Scientific Reports 9, 6397.

Wijnen H, Young MW. 2006. Interplay of circadian clocks and metabolic rhythms. Annual Review of Genetics 40, 409–448.

Xia T, Zhang L, Xu J, et al. 2017. The alternative splicing of EAM8 contributes to early flowering and short-season adaptation in a landrace barley from the Qinghai-Tibetan Plateau. Theor Appl Genet 130, 757–766.

Yan L, Fu D, Li C, et al. 2006. The wheat and barley vernalization gene VRN3 is an orthologue of FT. Proceedings of the National Academy of Sciences of the United States of America 103, 19581–19586.

Yan L, Loukoianov A, Blechl A, Tranquilli G, Ramakrishna W, SanMiguel P, Bennetzen JL, Echenique V, Dubcovsky J. 2004. The wheat VRN2 gene is a flowering repressor down-regulated by vernalization. Science 303, 1640–1644.

Yan L, Loukoianov A, Tranquilli G, Helguera M, Fahima T, Dubcovsky J. 2003. Positional cloning of the wheat vernalization gene VRN1. Proceedings of the National Academy of Sciences of the United States of America 100, 6263–6268.

Yu JW, Rubio V, Lee NY, et al. 2008. COP1 and ELF3 control circadian function and photoperiodic flowering by regulating GI stability. Molecular Cell 32, 617–630.

Zadoks JC, Chang; TT, Konzak CF. 1974. A decimal code for the growth stages of cereals. Weed research 14, 415–421.

Zagotta MT, Hicks KA, Jacobs CI, Young JC, Hangarter RP, Meeks-Wagner DR. 1996. The Arabidopsis ELF3 gene regulates vegetative photomorphogenesis and the photoperiodic induction of flowering. Plant Journal 10, 691–702.

Zakhrabekova S, Gough SP, Braumann I, et al. 2012. Induced mutations in circadian clock regulator Mat-a facilitated short-season adaptation and range extension in cultivated barley. Proceedings of the National Academy of Sciences of the United States of America 109, 4326–4331.

Zamir D. 2001. Improving plant breeding with exotic genetic libraries. Nature Review Genetics 2, 983–989.

Zheng B, Biddulph B, Li D, Kuchel H, Chapman S. 2013. Quantification of the effects of VRN1 and Ppd-D1 to predict spring wheat (Triticum aestivum) heading time across diverse environments. J Exp Bot 64, 3747–3761.

Zhu Z, Quint M, Anwer MU. 2022. Early Flowering 3 controls temperature responsiveness of the circadian clock independently of the evening complex. Journal of Experimental Botany 10.1093/jxb/erab473.

