## Supplementary Figures S1-S5 for "Novel exotic alleles of EARLY FLOWERING 3 determine plant development in barley"

### Figure S1. Weather data.

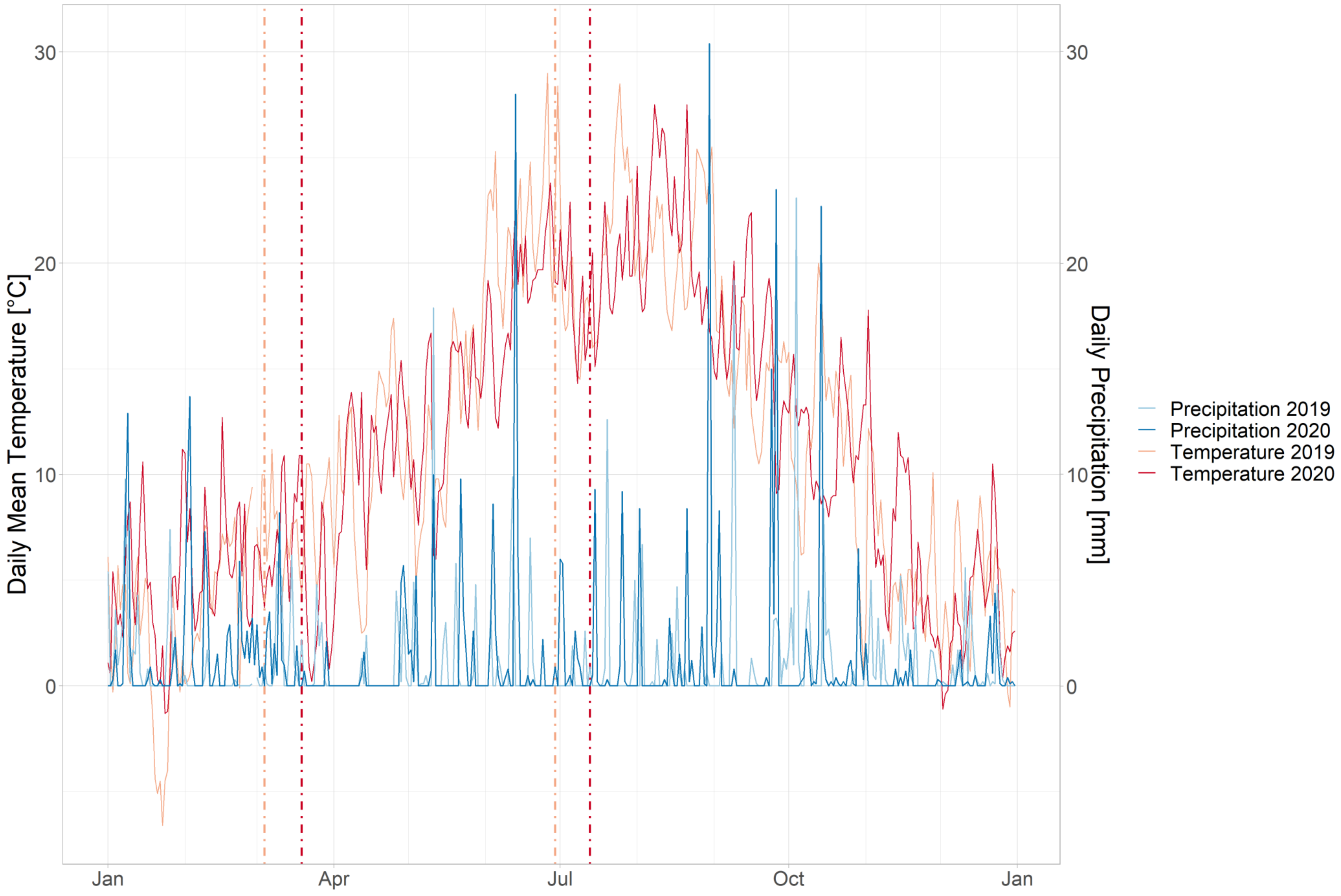


Temperature and Precipitation for both field trial years separately. Bright and dark red vertical lines show the respective vegetation period 2019 and 2020.

**Figure S2. Boxplots for all traits and both years separately.**

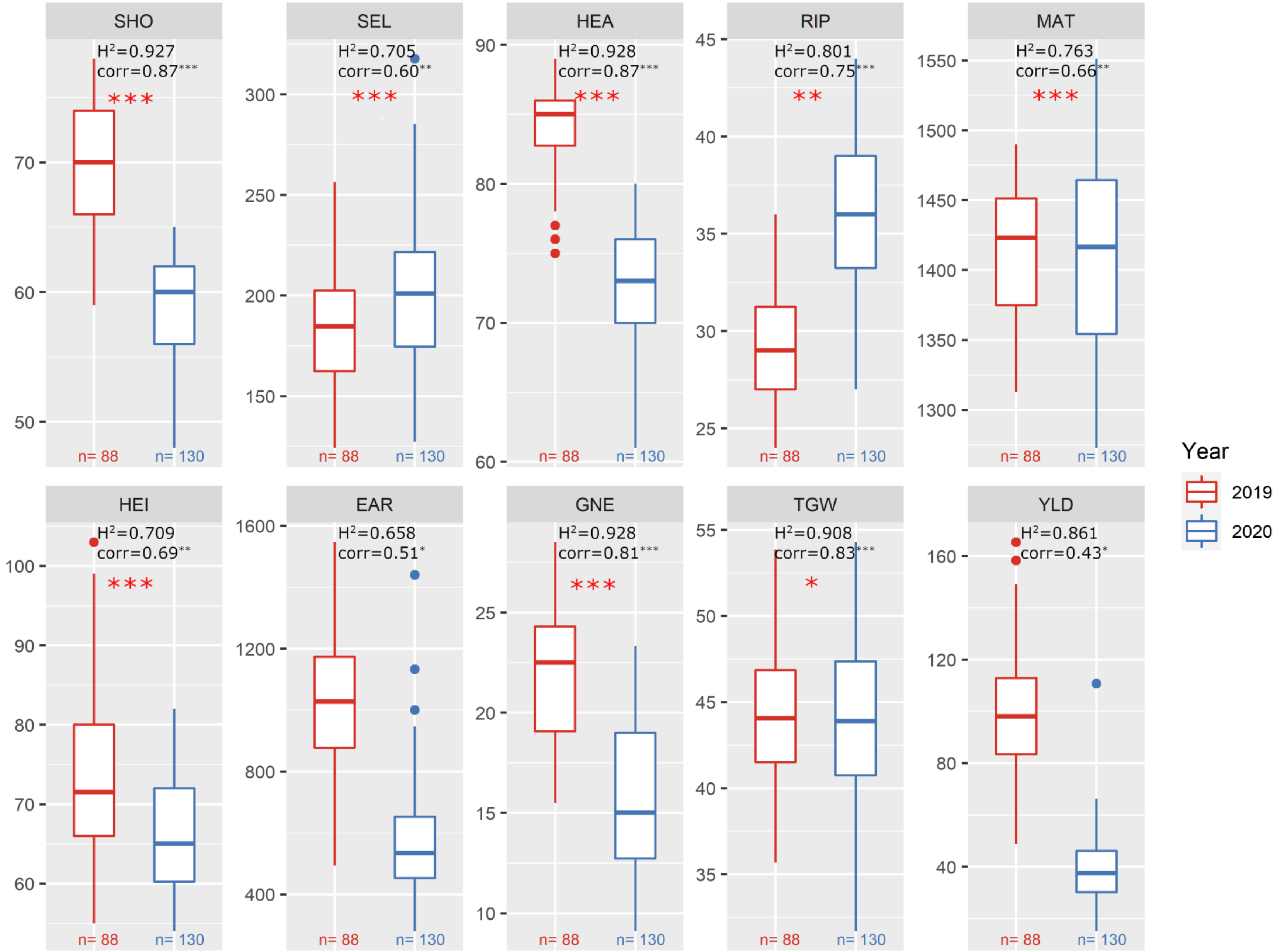


Comparison of all studied traits between 2019 and 2020 (red and blue boxplots, respectively). Units for each trait can be found in Table 1. Heritability ($H^2$) for traits and correlation (corr) of traits across years are noted. Red asterisks indicate a significant ANOVA for Line × Year interaction (*$p < 0.05$, ** $p < 0.01$ and *** $p < 0.001$).

### Figure S3A. Correlation of traits for 2019 and 2020 separately.

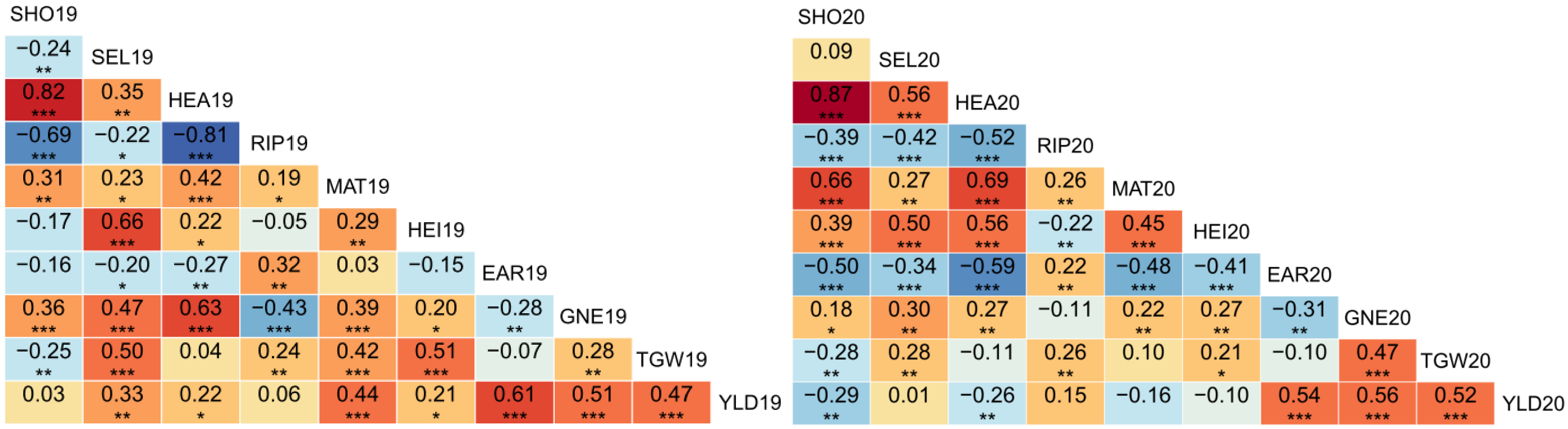


Pearson’s correlations between all traits for (A) all lines together and (B) *$ELF_{Hv}$* carrying lines (left) and *$ELF3_{Hsp}$* carrying lines (right) separately for 2019 and 2020. Asterisks indicate significant correlations between the respective traits (*p < 0.05, ** p < 0.01 and *** p < 0.001).

### Figure S3B. Correlation of traits for 2019 and 2020 separately.

$ELF_{Hv}$ carrying lines only

$ELF_{Hsp}$ carrying lines only

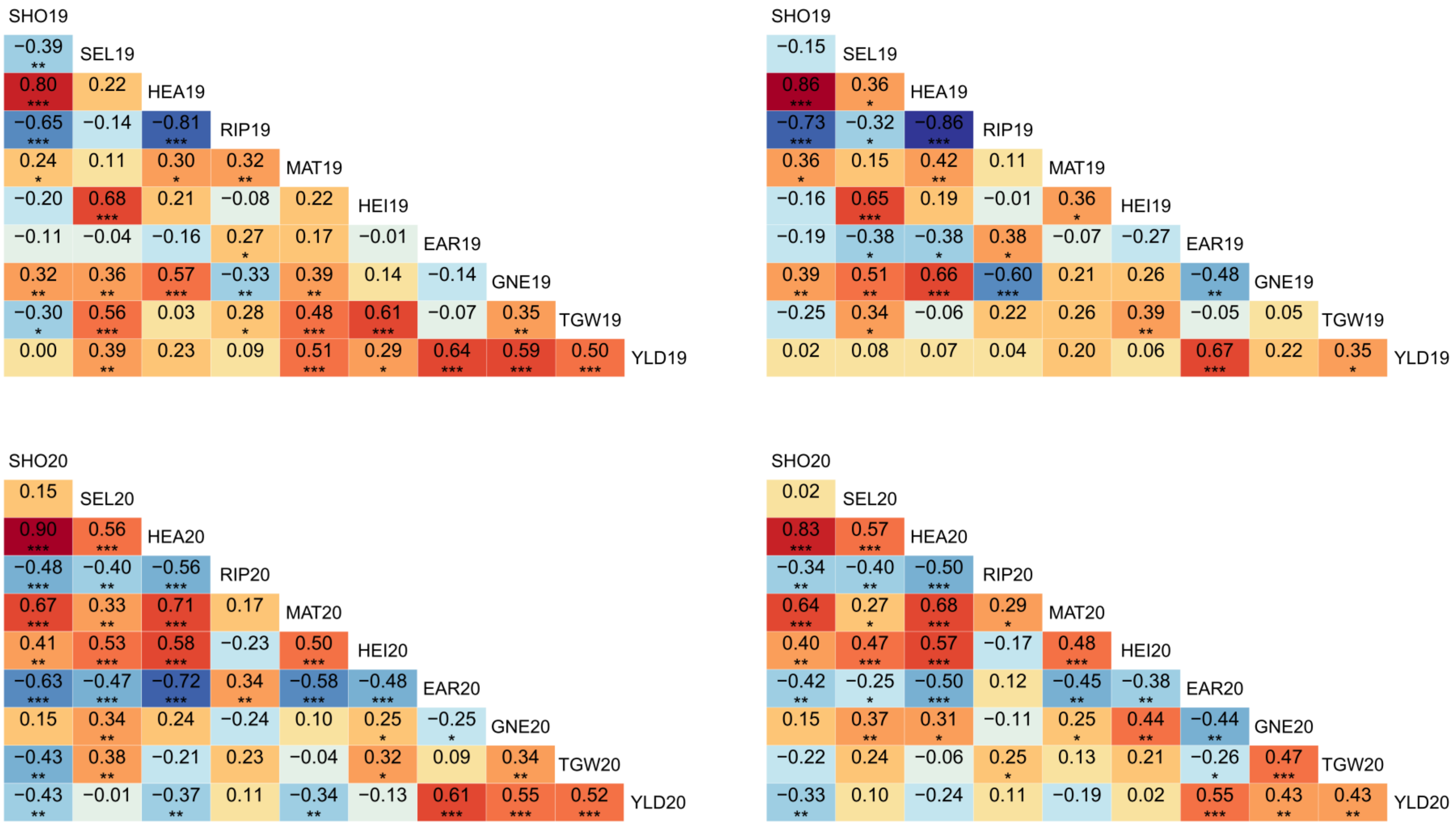


Pearson's correlations between all traits for (A) all lines together and (B) $ELF_{Hv}$ carrying lines (left) and $ELF3_{Hsp}$ carrying lines (right) separately for 2019 and 2020. Asterisks indicate significant correlations between the respective traits (*p < 0.05, ** p < 0.01 and *** p < 0.001).

**Figure S4. Trait differences between the two sister lines of each HIF pair ($ELF3_{Hsp}$ compared to $ELF3_{Hv}$) across years.**

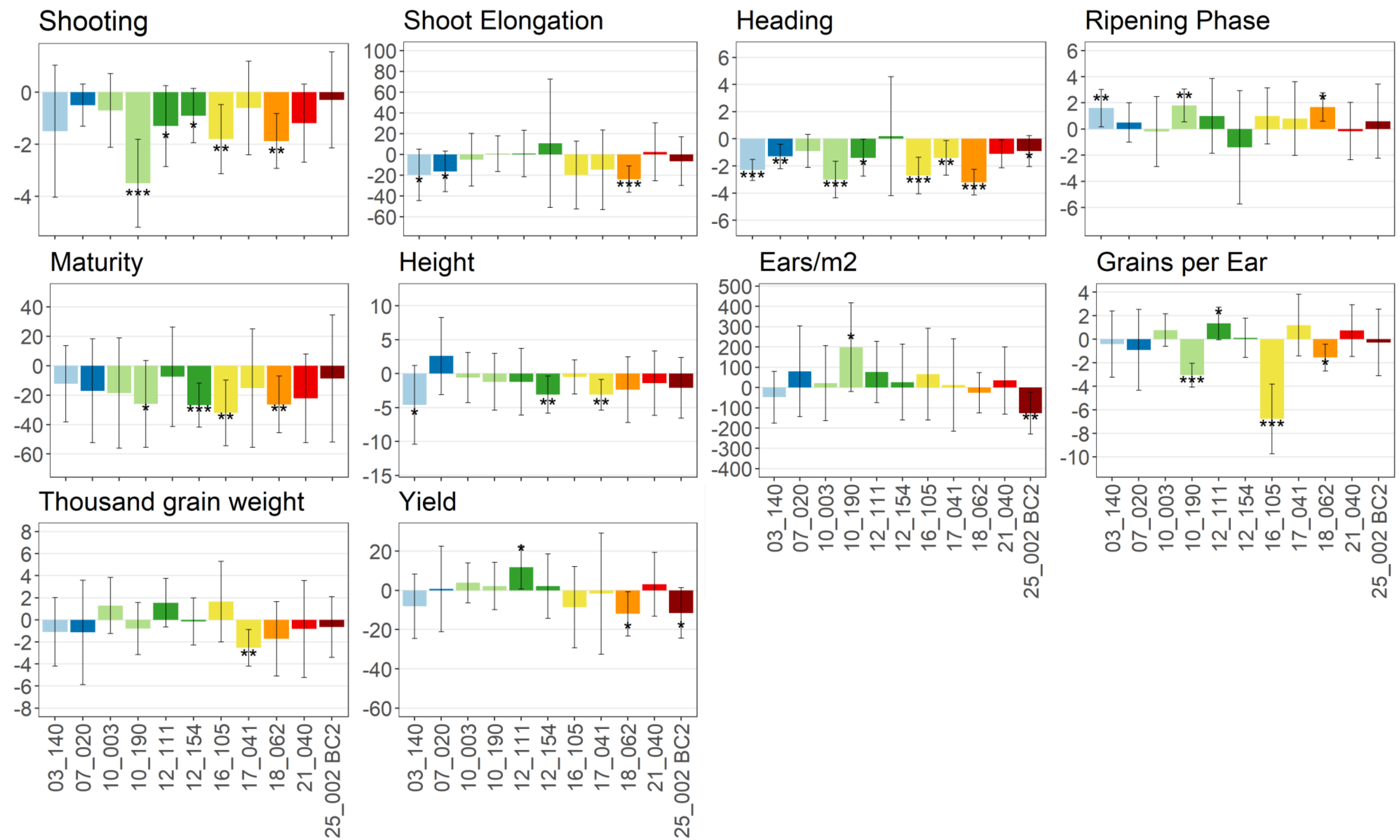


Lines with two identical first digits originate from the same wild donor. Trait units are given in Table 1. Asterisks indicate that the difference between sister lines is significantly different from zero (one-sample t-test, *p < 0.05, ** p < 0.01 and *** p < 0.001) and error bars show standard deviations. Mean differences and standard deviations are based on differences for each HIF pair per block. Columns are coloured depending on the ELF3 haplotype defined in Fig. 4C.

### Figure S5A. Segregating regions between HIF sister lines.

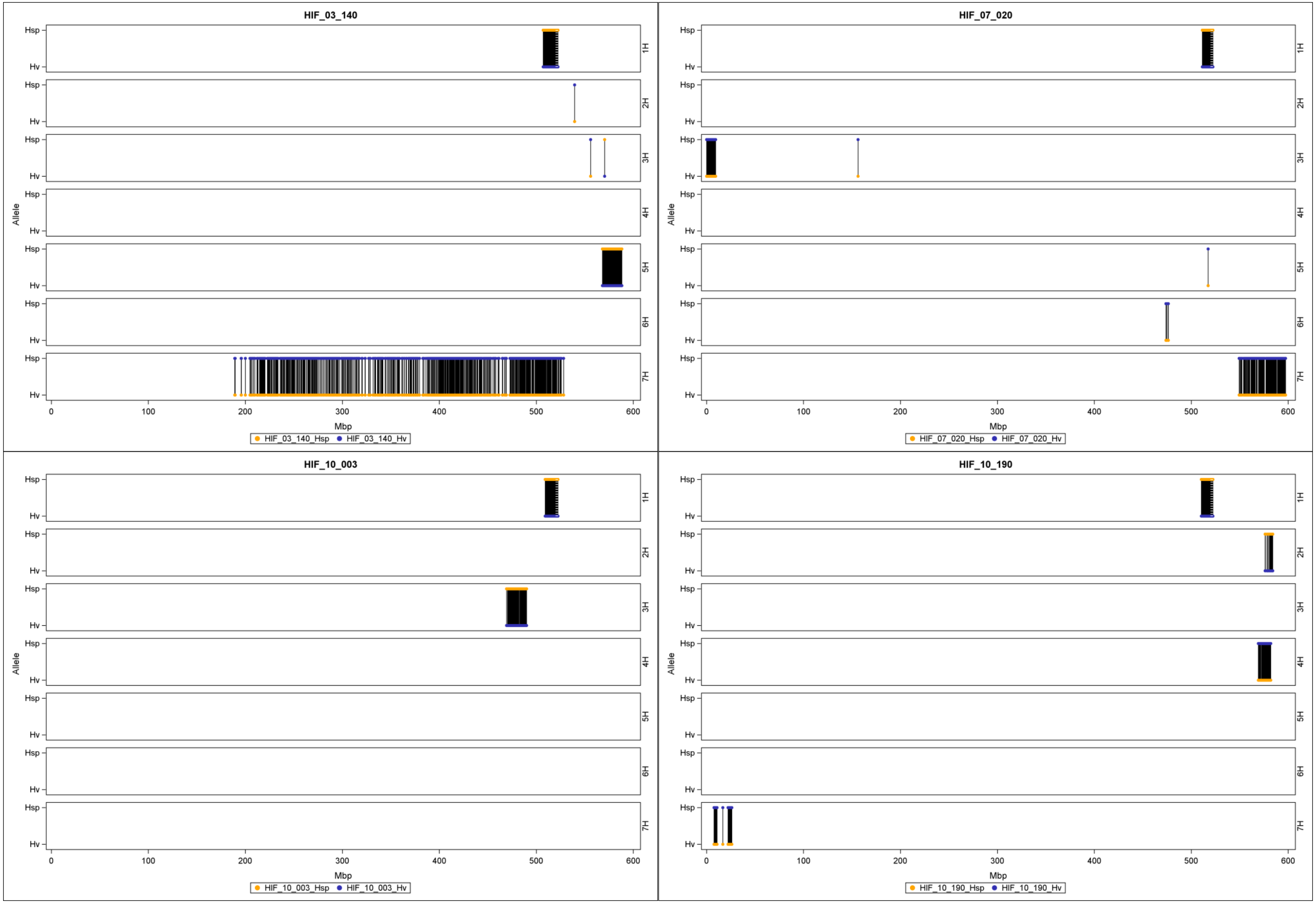


Comparison of the whole genome of all HIF lines based on the genotype data from the Infinium iSelect 50k SNP chip (Table S3). The dashed white line shows the *ELF3* position (derived from Morex reference sequence v2, Monat et al. (2019)) on chromosome 1H. Black bars represent differing regions between the $ELF3_{Hsp}$ (orange dots) and $ELF3_{Hv}$ (blue dots) sister lines with the upper and bottom line of each chromosome indicating the allele state (*Hsp* or *Hv*).

### Figure S5B. Segregating regions between HIF sister lines.

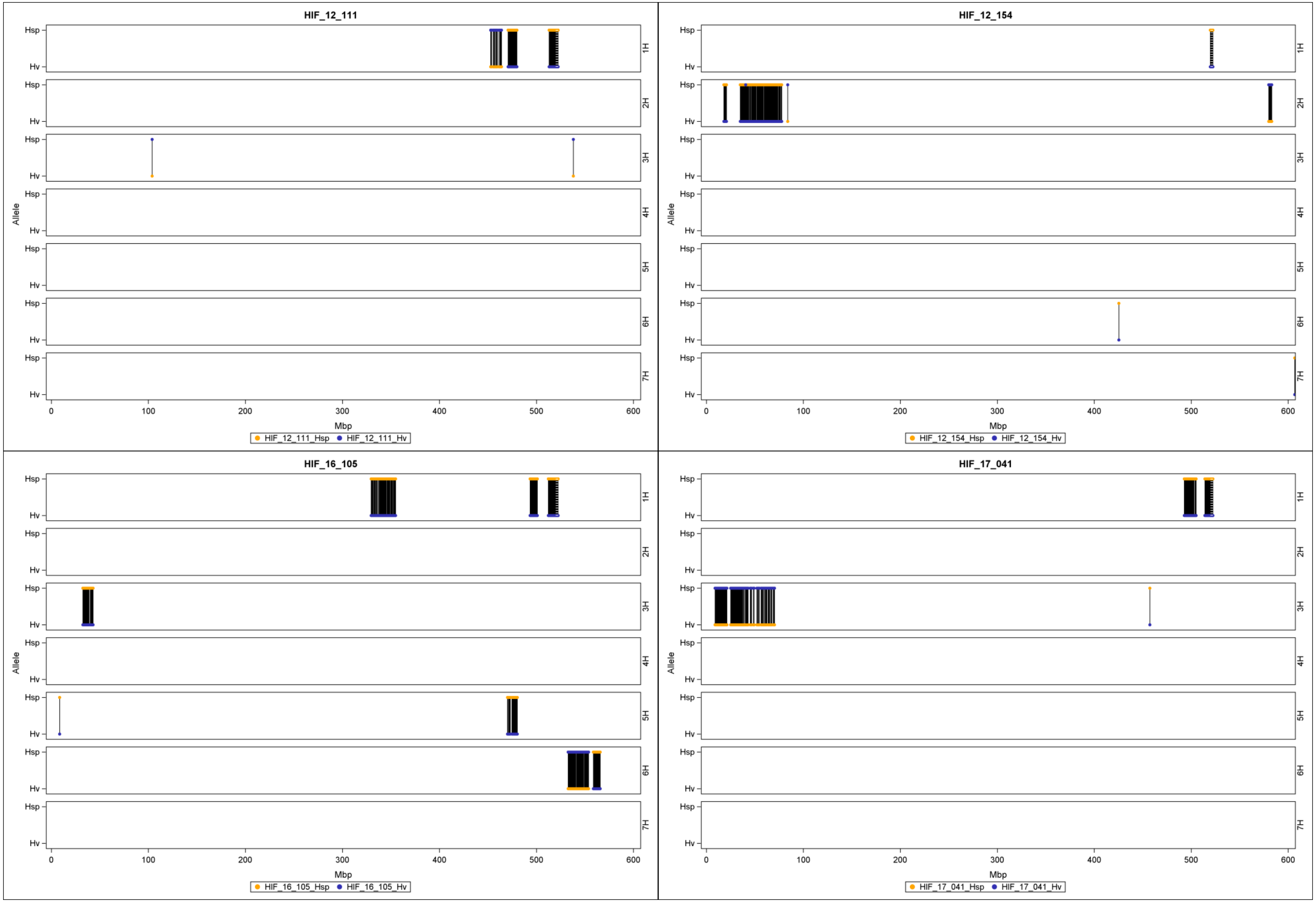


Comparison of the whole genome of all HIF lines based on the genotype data from the Infinium iSelect 50k SNP chip (Table S3). The dashed white line shows the *ELF3* position (derived from Morex reference sequence v2, Monat et al. (2019)) on chromosome 1H. Black bars represent differing regions between the $ELF3_{Hsp}$ (orange dots) and $ELF3_{Hv}$ (blue dots) sister lines with the upper and bottom line of each chromosome indicating the allele state (*Hsp* or *Hv*).

### Figure S5C. Segregating regions between HIF sister lines.

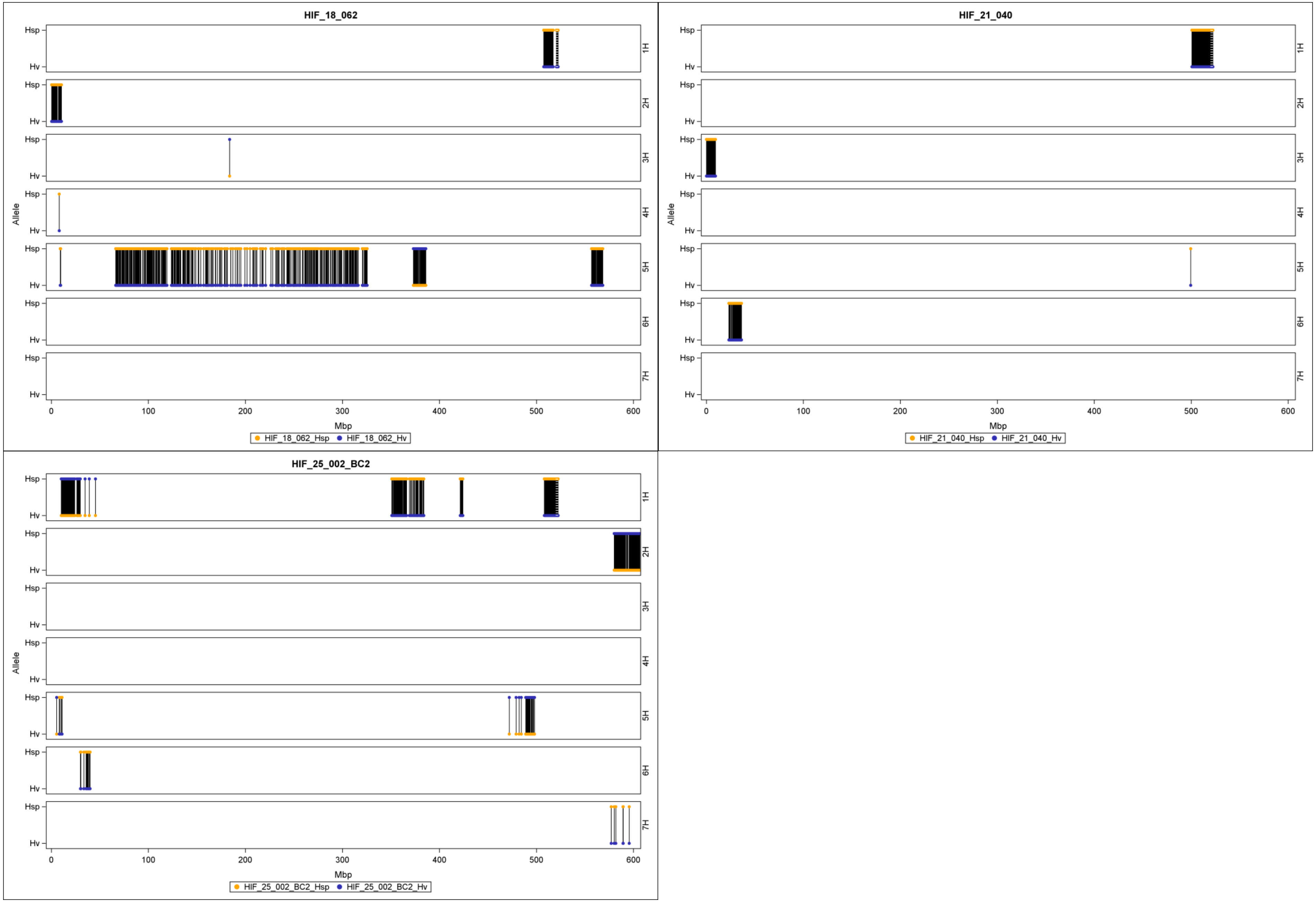


Comparison of the whole genome of all HIF lines based on the genotype data from the Infinium iSelect 50k SNP chip (Table S3). The dashed white line shows the *ELF3* position (derived from Morex reference sequence v2, Monat et al. (2019)) on chromosome 1H. Black bars represent differing regions between the $ELF3_{Hsp}$ (orange dots) and $ELF3_{Hv}$ (blue dots) sister lines with the upper and bottom line of each chromosome indicating the allele state (*Hsp* or *Hv*).
