## Supplementary Note S1 for "Novel exotic alleles of EARLY FLOWERING 3 determine plant development in barley"

*Repeatabilities and Heritabilities*

Calculation of repeatabilities (Rep) and heritabilities ($H^2$) was conducted for all HIF lines and Barke together (Table S5). For the developmental parameters, Rep and $H^2$ were calculated for both, days and GDD, revealing that for SEL and MAT, GDD exceeded days in both cases. Maurer *et al.* (2016) described, that GDD offers better estimates in these cases, as plant development and biochemical processes are temperature-dependent. Particularly, when comparing different years, it can be reasonable to use GDD rather than days. Days can outperform GDD when further temperature increase does not affect plant growth anymore or when the plant reaches a critical physiological stage. So, the best model for prediction depends on the respective developmental stage (McMaster, 1988). Consequently, for further analyses, GDD was used for the traits SEL and MAT and days was used for the remaining developmental traits. Lower $H^2$ values, for example for HEI and SEL, are in compliance with lower correlations between years (Figure S2), showing that they were strongly influenced by environmental conditions, emphasising the differences between the two trial years.

*Correlations*

Pearson's correlation coefficients (*r*) of traits were calculated between the two years (Figure S3) and also for both years separately (Figure S3). For single years, correlations are in compliance with previous findings in HEB-25 (Herzig *et al.*, 2018; Maurer *et al.*, 2016; Wiegmann *et al.*, 2019). With bigger plot sizes, correlations improved in 2020 for most traits, especially for EAR, since this is the most sensitive trait due to its measurement. Furthermore, regarding the relationship between Heading and Yield, correlations are stronger for the $ELF3_{Hv}$-carrying lines in both years (0.23 in 2019 and -0.37 with $p<0.01$ in 2020) compared to the $ELF3_{Hsp}$-carrying lines (0.07 in 2019 and -0.24 in 2020), showing that the effects are quite dynamic and the environment effects are possibly dampened by the $ELF3_{Hsp}$-Allele. Altogether, results differ between years which indicates that both field years are rather different from each other and it is reasonable to evaluate both years separately.

Cross-correlations of traits between years were high (0.51-0.87) and highly significant, except for the trait YLD (0.43), showing that the environmental effect is highest for this trait. SHO, HEA, RIP, GNE and TGW show very high correlations ($r > 0.7$), indicating that these traits were less influenced by environmental cues than the remaining traits. However, this is actually in contradiction to the high $H^2$ (Table S5) estimates for YLD.

*Impacts of the environment on phenotypes*

At higher temperatures, as in 2020 (Supplementary Table S4), development of plants is accelerated and due to the lack of time for assimilation and grain filling, the recorded values of the yield components might be lower. Precipitation, or rather the lack of it, also determines grain filling and maturation. Possibly, early drought stress in 2020 and late rainfalls compared to evenly distributed rainfall in 2019 (Supplementary Table S4) may therefore also have resulted in lower yield component values. Day length not only influences the amount of photosynthesis, but just like temperature, also the control of plant development by the circadian clock (Bendix *et al.*, 2015; Calixto *et al.*, 2015; Harmer, 2009; Nusinow *et al.*, 2011; Wijnen and Young, 2006). Since the vegetation period started two weeks later in 2020, days were longer than in 2019 for the major part of the growing season (Supplementary Table S4), which possibly also led to a faster development in 2020. Accordingly, cumulative day lengths were higher for developmental traits in 2019. Furthermore, $ELF3_{Hsp}$-carrying lines needed less photoperiod to reach the respective developmental stages compared to $ELF3_{Hv}$-carrying lines in both years (Supplementary Table S7) which could be due to the role of *ELF3* in the circadian clock controlling plant development.

Finally, the found effects are probably due to an interaction of many factors, including temperature, precipitation and day length. Higher temperature, less rain in the beginning of the vegetation period and greater day lengths might all have led to a faster development and lower yield components in 2020 and possibly to more and different $ELF3_{Hsp}$-effect differences in 2020. However, as already mentioned, a limiting factor of this study is the lack of optimal conditions, e.g. too much variation in temperature (and only daily average temperature whereas daily maximum temperature would also be interesting), precipitation (possibly also at some points even a lack of precipitation) and photoperiod. All these facts also support the decision for an annual evaluation.
